# CD90 identifies distinct fractions of muscle stem cells with different modalities of activation and quiescence maintenance

**DOI:** 10.1101/2024.10.25.619970

**Authors:** Eyemen Kheir, Michela Libergoli, Samuele Metti, Jonas Brorson Jensen, Jakob Wang, Niels Jessen, Francesca Florio, Francesca Murganti, Serena Di Savino, Marzia Belicchi, Chiara Villa, Kristian Vissing, Mayank Verma, Lorenzo Giordani, Paolo Bonaldo, Yvan Torrente, Jean Farup, Stefano Biressi

## Abstract

Stem cell transition from quiescence to activation is crucial to guarantee productive tissue regeneration. Here we show that CD90 diversifies quiescent muscle stem cells (MuSCs) in murine and human muscle into two subpopulations differing in the kinetics of activation, CD90^+ve^ MuSCs exhibiting a faster exit quiescence and predominating the initial phases of regeneration compared to CD90^-ve^ MuSCs. In the absence of injury, the CD90^+ve^ fraction is primed toward activation through an active CD90-AMPK axis but is maintained in quiescence through signals from the extracellular matrix. Our studies show that Collagen VI, which is preferentially expressed by CD90^+ve^ MuSCs, binds to the Calcitonin receptor and plays a role in this context. Moreover, while the number of CD90^+ve^ and CD90^-ve^ subpopulations is similar in healthy muscles, the CD90^-ve^ fraction predominates in the muscles of murine models of Duchenne and Ullrich congenital muscular dystrophies. These findings provide novel insights into the mechanistic determinants of MuSCs functional heterogeneity and have implications for understanding the stimulation of repair in dystrophic muscle.

## INTRODUCTION

Skeletal muscle satellite cells (MuSCs) represent a muscle-specific stem cell population characterized by the expression of Pax7, which govern skeletal muscle repair (Scharner and Zammit, 2011). In homeostatic conditions, MuSCs reside in a quiescent state, where they are poised for activation (Montarras et al., 2013; van Velthoven and Rando, 2019). In response to muscle injury, MuSCs become mitotically active, differentiate and self-renew to regenerate the tissue (Scharner and Zammit, 2011). Muscle regeneration requires the orchestrated contribution of a heterogeneous group of support cells, which include fibro-adipogenic progenitors (FAPs) and endothelial cells (Wosczyna and Rando, 2018). In response to distal trauma, MuSCs enter a state of ‘partial’ activation named G_Alert_ characterized by increased cell size and mitochondrial content (Rodgers et al., 2014). This has been proposed as a mechanism that MuSCs use to preempt the response to damage (Rodgers et al., 2014). There is much principal and clinical interest in understanding the mechanisms regulating quiescence and activation, as alterations of this balance may result in failure of muscle regeneration (Baghdadi et al., 2018; Bjornson et al., 2012; de Morrée et al., 2017; Hausburg et al., 2015; Jash et al., 2014; Machado et al., 2018; Mourikis et al., 2012; van Velthoven et al., 2017).

An increasing number of studies collectively indicated that the pool of MuSCs is more heterogeneous than initially contended (Biressi and Rando, 2010; Tierney and Sacco, 2016). However, among the numerous studies showing molecular heterogeneity in myogenic precursor cells, only in few cases have these molecular differences been linked to functional heterogeneities (Barruet et al., 2020; Beauchamp et al., 2000; Chakkalakal et al., 2014; De Micheli et al., 2020; Dell’Orso et al., 2019; Der Vartanian et al., 2019; García-Prat et al., 2020; Kuang et al., 2007; Tanaka et al., 2009). We report here that the GPI-anchored protein CD90 (Thy-1) splits the pool of MuSCs into two subpopulations. CD90 is reportedly expressed in different stem cell populations, including mesenchymal and hematopoietic stem cells (Herrera-Molina et al., 2013). Our findings identify CD90 as a novel marker of MuSCs heterogeneity and establish its role as a promoter of MuSCs activation and proliferation. Intriguingly, in the absence of trauma, CD90^+ve^ MuSCs present a population-specific molecular program, which involves the differential expression of Collagen VI (Col6) and its binding to the calcitonin receptor, thus ensuring quiescence by restraining the propensity of CD90^+ve^ MuSCs toward activation. We also observed that the depletion of CD90^+ve^ cells in regenerating muscle delays regeneration and that the balance in abundance between CD90^+ve^ and CD90^-ve^ MuSCs is altered in animal models of different forms of muscular dystrophy. These studies extend the concept of stem cell diversification to the processes controlling the transition from quiescence to proliferation and increase our knowledge of the cellular dynamics paralleling defective regeneration in dystrophic muscle.

## RESULTS

### CD90 marks distinct fractions of quiescent MuSCs

We have previously reported that CD90 marks a fraction of FAPs associated with fibrotic remodeling in diabetic patients (Farup et al., 2021). Here, we fractionated mouse skeletal muscle into various subpopulations of muscle-resident cells and stained them with an anti-CD90 antibody for FACS analysis. CD90 was diversifying not only the FAPs, but also the pool of MuSCs into two fractions (**Figures 1A-C**). The heterogeneous expression of CD90 in MuSCs was confirmed by CyTOF investigation of a publicly available dataset (Porpiglia et al., 2017) (**Figure 1D**). We further corroborated these observations by using *Pax7^CreERT2^;R26R^EYFP^*and *Myf5^CreER^;R26R^EYFP^* transgenic mice, in which the expression of Cre recombinase from the *Pax7* and *Myf5* loci, respectively, induces the YFP reporter in MuSCs upon tamoxifen injection (Biressi et al., 2013; Murphy et al., 2011). In these strains approximately half of the YFP^+ve^ MuSCs were CD90^+ve^, similar to those from wild-type mice (**Figures 1E**, **S1A**, and **S1B**). We validated the specificity of CD90 staining by using two different clones of anti-CD90 antibody (**Figures S1A**). Two variants of CD90, CD90.1 and CD90.2, exist in different mouse strains (Herrera-Molina et al., 2013). The antibodies we employed recognize the CD90.2 variant, expressed in the *C57BL/6J* background of the animals we have used so far. To further demonstrate the specificity of CD90 staining, we backcrossed *Myf5^CreER^;R26R^EYFP^* mice in the *FVB* background, which produces the CD90.1 variant. The absence of staining in the FVB MuSCs confirmed that the CD90 signal in the *C57BL/6J* background is specific (**Figure S1B**). We confirmed the myogenic identity of CD90^+ve^ and CD90^-ve^ MuSCs by staining them with an anti-Pax7 antibody (**Figure S1C**). Moreover, CD90^+ve^ and CD90^-ve^ MuSCs formed Myogenin (Mgn)^+ve^ myotubes *ex vivo* (**Figure 1F**).

**Figure 1.**
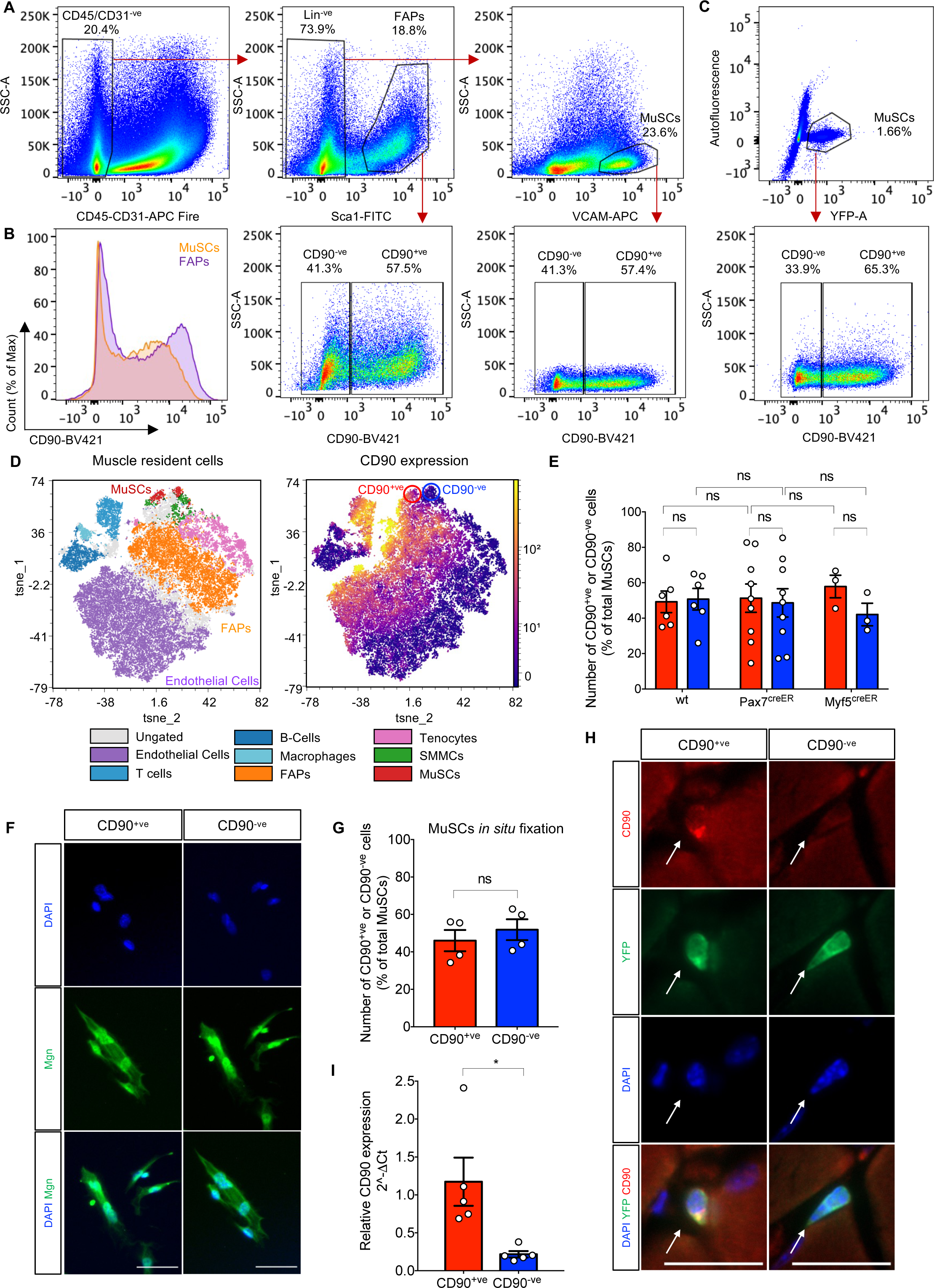
CD90 identifies two distinct subpopulations of quiescent MuSCs. (**A**) FACS profiles of FAPs and MuSCs from lower hindlimb muscles of uninjured *C57/BL6J* mice stained with antibodies against CD45, CD31, Sca1, VCAM, and CD90. FAPs are identified as CD45^-ve^/CD31^-ve^/Sca1^+ve^, MuSCs as Lin^-ve^ (CD45^-ve^/CD31^-ve^/Sca1^-ve^)/VCAM^+ve^. (**B**) FACS histogram showing CD90 expression in MuSCs and FAPs, identified as in (A). (**C**) FACS profile of YFP^+ve^ MuSCs of *Pax7^CreERT2/wt^;R26R^EYFP/wt^*mice stained with an antibody against CD90. (**D**) t-Stochastic neighbor embedding (tSNE) plots showing the distribution of muscle-resident populations (left; color codes for each cell population are indicated below the panel) and the corresponding expression pattern of CD90 (right; cells are color-coded according to CD90 intensity) in uninjured murine muscle analyzed by CyTOF (Porpiglia et al, 2017). Red and blue circles enclose CD90^+ve^ and CD90^-ve^ MuSCs, respectively. (**E**) Quantification of CD90 heterogeneity in MuSCs from hindlimb muscles identified as Lin^-ve^/VCAM^+ve^ cells in *C57/BL6J* (wt) mice, and as YFP^+ve^ cells in *Pax7^CreERT2/wt^;R26R^EYFP/wt^*and *Myf5^CreERT2/wt^;R26R^EYFP/wt^* mice (respectively indicated as Pax7^CreER^ and Myf5^CreER^). Error bars represent mean ± SEM, n≥3. Two-way ANOVA test performed. (**F**) Representative images of Mgn immunostaining in CD90^+ve^ and CD90^-ve^ MuSCs isolated from uninjured hindlimb muscles and cultured for 6 days. DAPI stains the nuclei. Scale bar: 50 μm. (**G**) Quantification of CD90^+ve^ and CD90^-ve^ *in situ* fixed YFP^+ve^ MuSCs. Error bars represent mean ± SEM, n=4. (**H**) Immunofluorescence analysis of CD90 in *gastrocnemius* muscle sections of *Pax7^CreERT2/wt^;R26R^EYFP/wt^* mice. Anti-YFP antibodies and DAPI were used to identify MuSCs and nuclei. Arrows indicate MuSCs. Scale bar: 20 μm. (**I**) RT-qPCR analysis of CD90 relative expression in CD90^+ve^ and CD90^-ve^ MuSCs isolated from uninjured hindlimb muscles of *Pax7^CreERT2/wt^;R26R^EYFP/wt^* mice. Error bars represent mean ± SEM, n=5.

The enzymatic digestion required to isolate MuSCs might perturb their quiescence state (Machado et al., 2018; van Velthoven et al., 2017). To investigate whether truly quiescent MuSCs heterogeneously express CD90, we performed an *in situ* fixation protocol before cell dissociation that allows capturing MuSCs in their native quiescent state. FACS analysis confirmed that CD90 is expressed by half of the MuSCs (**Figures 1G** and **S1D**). The presence of CD90^+ve^ and CD90^-ve^ MuSCs in the quiescent muscle tissue *in situ* was confirmed in *gastrocnemius* muscle cryosections and myofibers isolated from the *soleus* muscle (**Figures 1H** and **S1E**).

RT-qPCR analysis revealed that CD90^+ve^ and CD90^-ve^ MuSCs express dissimilar levels of CD90, indicating a transcriptional control of CD90 diversification (**Figure 1I**). RT-qPCR and protein analysis revealed that that CD90 expression does not match with previously reported heterogeneity markers (**Figures S2A-E**)(Beauchamp et al., 2000; Cerletti et al., 2008; Chakkalakal et al., 2014; Der Vartanian et al., 2019; García-Prat et al., 2020; Kuang et al., 2007; Rocheteau et al., 2012). Overall, our data show that quiescent MuSCs can be divided into two subpopulations that are proportionally similar but characterized by different levels of CD90.

### CD90^+ve^ and CD90^-ve^ MuSCs have different propensity to activate and enter G_Alert_ state

Using tamoxifen-injected *Pax7^CreERT2^;R26R^EYFP^*mice to identify MuSCs, we characterized the activation and proliferation dynamics of CD90^+ve^ and CD90^-ve^ MuSCs by employing three experimental paradigms (**Figure 2A**). Firstly, we FACS-purified YFP^+ve^/CD90^+ve^ and YFP^+ve^/CD90^-ve^ MuSCs from uninjured muscles and evaluated their activation before the first cell division (Rodgers et al., 2014). CD90^+ve^ MuSCs showed an increase in activation markers, such as an increment in cell size and in MyoD protein accumulation, as well as in the intensity of the YFP signal, a read-out of the *Rosa26* locus activity; previously used as an indicator of MuSCs activity (**Figures 2B-D**, and **S3A-D**)(Boonsanay et al., 2016; de Morrée et al., 2017; Hausburg et al., 2015; Rodgers et al., 2014). Given their higher propensity to activate, we reasoned that CD90^+ve^ MuSCs would also be more proliferative. Indeed, when we analyzed the sorted MuSCs subpopulations at a later time-point (i.e., 2.5 days *in vitro*), we observed that the increased MyoD expression parallels an accelerated entry into the cell cycle in CD90^+ve^ MuSCs (**Figure S3E-F**). Intriguingly, experiments in which CD90^+ve^ and CD90^-ve^ MuSCs were co-cultured suggest that sharing the same medium does not significantly influence these dynamics (**Figure S3E-F**). Secondly, we employed an *in vivo* needle-injury model characterized by a robust regenerative response (**Figure S3G** and **S3H**). At the earliest time-point after muscle injury, CD90^+ve^ MuSCs exhibited a significant increase in cell size and YFP reporter activity compared to the CD90^-ve^ counterpart (**Figures 2E-F**). In keeping with these observations, a significantly higher fraction of CD90^+ve^ MuSCs incorporated EdU in vivo at 1.5, 2.75, and 6 days after injury compared to its CD90^-ve^ counterpart (**Figure 2G**). Finally, we considered the transition from quiescence to the G_Alert_ state, which is induced in MuSCs by distal injury (Rodgers et al., 2014). Despite both subpopulations of MuSCs becoming progressively “alerted”, the CD90^+ve^ MuSCs exhibited faster G_0_ to G_Alert_ transition, shown by the increase in cell size, YFP intensity, and mitochondria content (**Figures 2H-M**). These findings suggest that, in the pool of quiescent MuSCs, it is possible to identify a CD90^+ve^ fraction, which is primed for rapid activation, and a more dormant CD90^-ve^ fraction.

**Figure 2.**
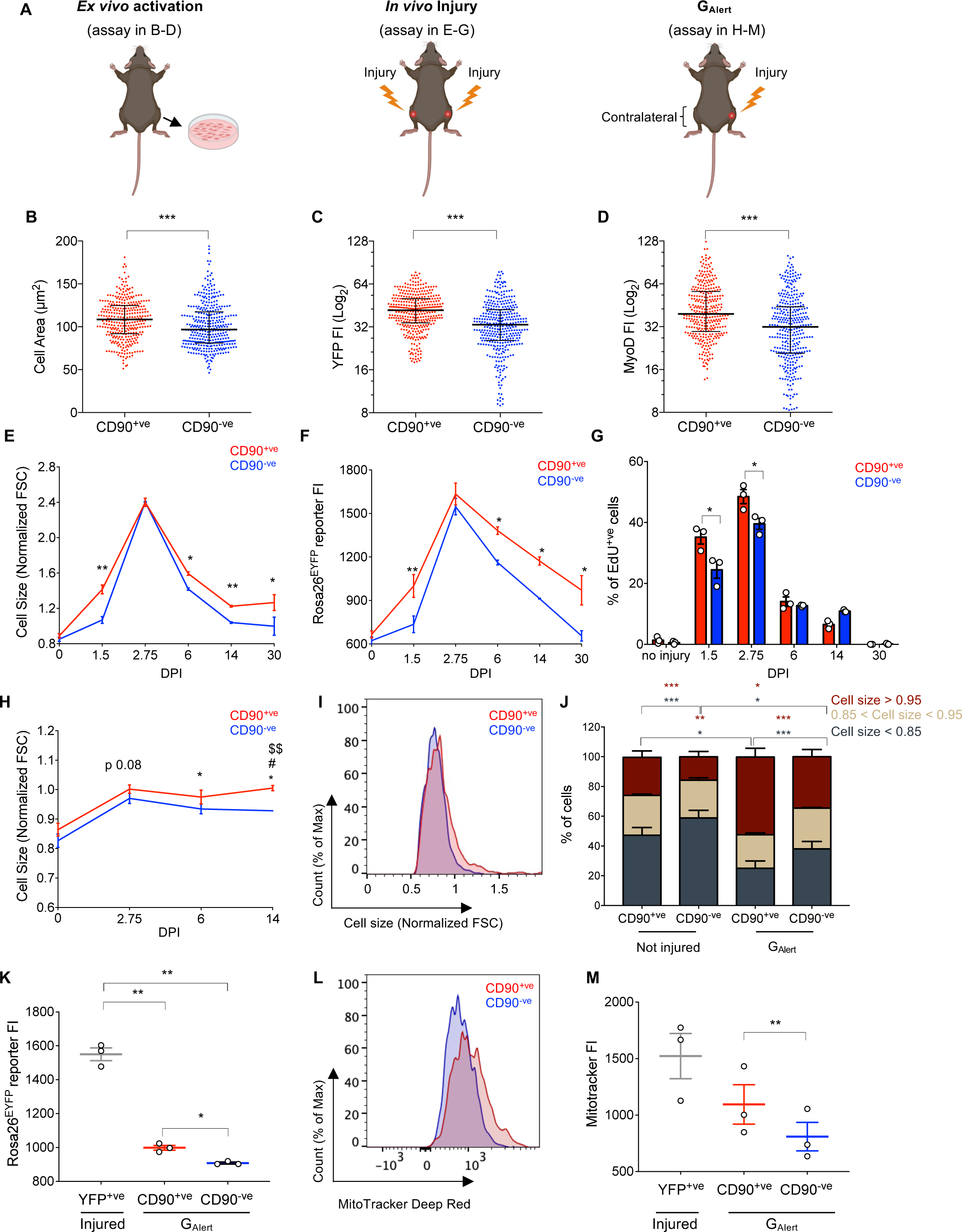
CD90^+ve^ and CD90^-ve^ MuSCs have different propensity to activate and enter G_Alert_ state. (**A**) Schematic representation of the MuSCs activation experiments. (**B-D**) Measurement of cell area (B), YFP (*Rosa26* locus activity) (C) and MyoD (D) fluorescence intensity (FI) in CD90^+ve^ and CD90^-ve^ MuSCs FACS-isolated from uninjured hindlimb muscles of *Pax7^CreERT2/wt^;R26R^EYFP/wt^* mice after 16 hours in culture. See Figure S3A-C for two additional independent experiments. Each dot represents a cell. Error bars represent median with interquartile range; the number of cells is ≥ 300. A two-tailed Mann-Whitney test was performed. (**E**) Measurements of FSC-A by FACS of CD90^+ve^ and CD90^-ve^ MuSCs from hindlimb muscles of *Pax7^CreERT2/wt^;R26R^EYFP/wt^* mice at 0 (no injury), 1.5, 2.75, 6, 14 and 30-days post-injury (DPI). To assess cell size, FSC-A values were normalized by the FSC-A of 15 μm standard beads. Error bars represent mean ± SEM, n=3. (**F**) FACS analysis of YFP fluorescence intensity showing that CD90^+ve^ MuSCs from hindlimb muscles of *Pax7^CreERT2/wt^;R26R^EYFP/wt^* mice display an increased YFP expression regulated by the *Rosa26* promoter at specified time-points of muscle injury. Data are expressed as median fluorescence intensity (FI) of both subpopulations. Error bars represent mean ± SEM, n=3 for each time-point. (**G**) Percentage of EdU^+ve^ MuSCs after injury. MuSCs were identified as YFP^+ve^ cells from hindlimb muscles of *Pax7^CreERT2/wt^;R26R^EYFP/wt^* mice. EdU was administered 12 hours before sacrifice. Error bars represent mean ± SEM, n=3 for each time-point. (**H**) Measurements of FSC-A of CD90^+ve^ and CD90^-ve^ MuSCs in contralateral hindlimb muscles at 0 (no injury), 2.75, 6, and 14 DPI. MuSCs were identified as YFP^+ve^ cells from *Pax7^CreERT2/wt^;R26R^EYFP/wt^* mice. To assess cell size, FSC-A was normalized by FSC-A of 15 μm standard beads. * refers to comparisons between CD90^+ve^ and CD90^-ve^ MuSCs; $ to comparison of CD90^+ve^ MuSCs at 0 *versus* 14 DPI; # to comparison of CD90^-ve^ MuSCs at 0 *versus* 14 DPI. Error bars represent mean ± SEM, n=3 for each time-point. A one-way Anova was used for multiple comparisons. (**I**) FACS histogram showing cell size distribution of the two subpopulations of MuSCs in contralateral muscles of injured mice at 14 days post-injury (DPI). FSC-A was normalized by FSC-A of 15 μm standard beads. (**J**) Quantification of the percentage of CD90^+ve^ and CD90^-ve^ MuSCs with indicated cell size ranges normalized by FSC-A of 15 μm standard beads in not injured and contralateral muscles of injured mice at 14 days post-injury (DPI). Error bars represent mean ± SEM, n=5. (**K**) FACS analysis of the YFP median fluorescence intensity (FI) of YFP^+ve^/CD90^+ve^ and YFP^+ve^/CD90^-ve^ MuSCs in contralateral muscles of *Pax7^CreERT2/wt^;R26R^EYFP/wt^* mice at 7 days post-injury (DPI). YFP intensity of total YFP^+ve^ MuSCs from injured muscles at 7 DPI is also represented. Error bars represent mean ± SEM, n=3. One-way ANOVA test was performed. (**L**, **M**) Representative FACS histograms of active mitochondria in CD90^+ve^ and CD90^-ve^ fractions of MuSCs (identified as YFP^+ve^ in *Pax7^CreERT2/wt^;R26R^EYFP/wt^* mice) in contralateral muscles 6 days after injury (L) and quantification (M). The median fluorescence intensity (FI) of MitoTracker Deep Red staining was used as a read-out of mitochondrial activity. The intensity of MuSCs from injured muscles is also represented. Error bars represent mean ± SEM, n=3. A one-way ANOVA test was performed.

### The fate of CD90^+ve^ MuSCs during muscle repair

In keeping with the accelerated activation of CD90^+ve^ MuSCs, we investigated whether the proportion of the two subpopulations would change during regeneration (**Figures 1E** and **3A**). We found that the CD90^+ve^ fraction predominates during the initial phase of regeneration. However, when the regeneration is complete, the proportion of the two subpopulations returned to pre-injury values (**Figure 3A**). This observation encouraged us to further investigate the cellular dynamics controlling the fate of the CD90^+ve^ MuSCs.

**Figure 3.**
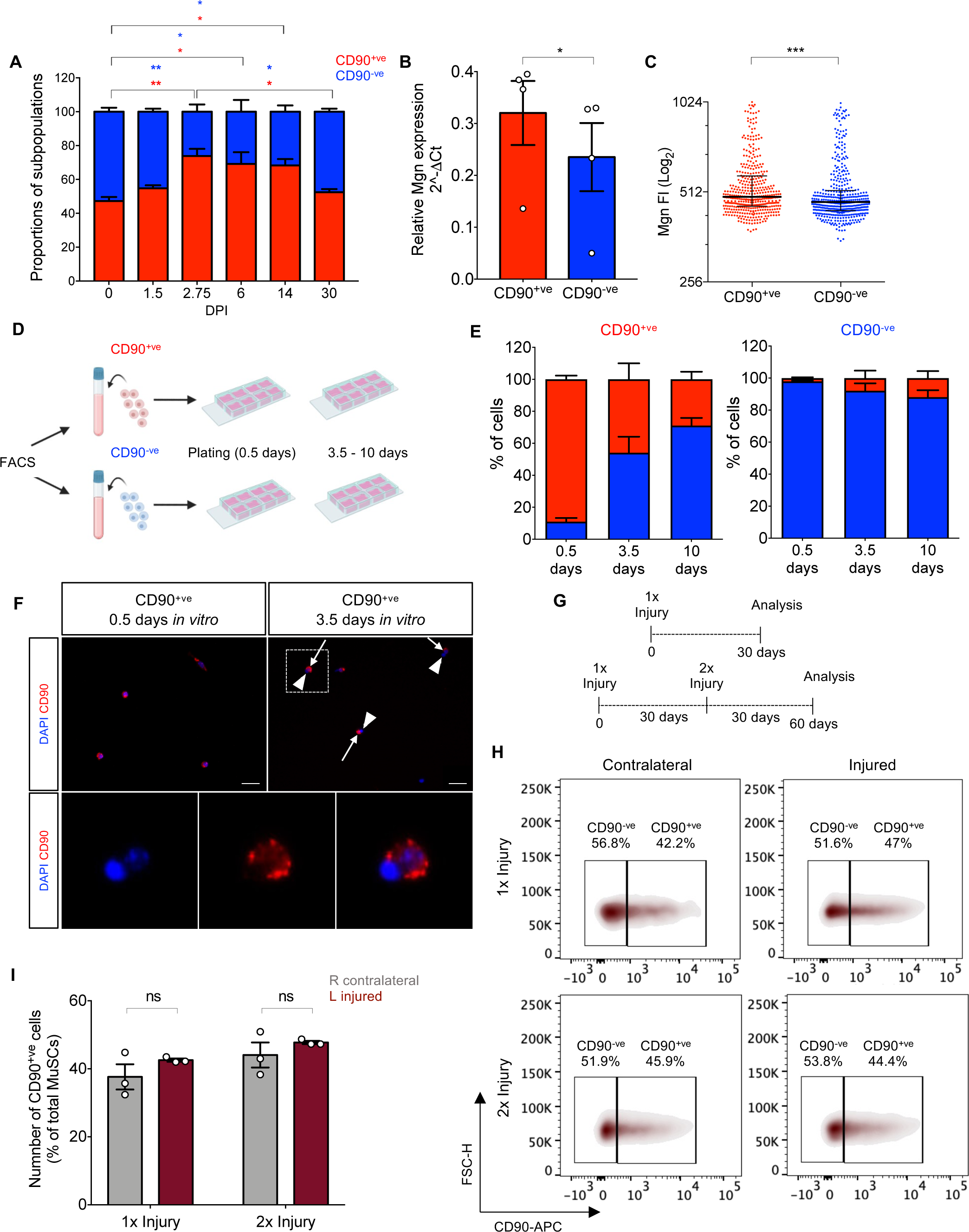
The fate of CD90^+ve^ MuSCs during muscle repair. (**A**) The proportion of CD90^+ve^ and CD90^-ve^ MuSCs in injured hindlimb muscles at specified time-points of muscle injury (DPI) was quantified, normalized with values obtained in uninjured contralateral muscles, and expressed as a percentage. Error bars represent mean ± SEM, n=3 for each time-point. Two-way ANOVA test was performed. Asterisks refer to comparisons between the specified time-points for CD90^+ve^ (red asterisks) and CD90^-ve^ (blue asterisks) MuSCs. (**B**) RT-qPCR analysis of Mgn relative gene expression in CD90^+ve^ and CD90^-ve^ MuSCs isolated 7 days post-injury. Error bars represent mean ± SEM, n=4. (**C**) Representative immunofluorescence analysis of Mgn fluorescence intensity (FI) of CD90^+ve^ and CD90^-ve^ MuSCs isolated at 7 days post-injury. Each dot represents a cell. Error bars represent median with interquartile range, the number of cells/replicate is ≥ 400. A two-tailed Mann-Whitney test was performed. See Figure S4A for two additional independent experiments. (**D**) Schematic representation of lineage study experiments shown in (E). (**E**) Quantification of the percentage of CD90^+ve^ and CD90^-ve^ cells within the CD90^+ve^ and CD90^-ve^ YFP^+ve^ populations sorted from uninjured tamoxifen-injected *Pax7^CreERT2^;R26R^EYFP^* mice and analyzed at 0.5-, 3.5- and 10-days post-plating. Error bars represent mean ± SEM, n=3. (**F**) CD90 immunofluorescence in CD90^+ve^ MuSCs cultures at 0.5- and 3.5-days post-plating. Arrows and arrowheads indicate newly formed daughter cells that are positive or negative, respectively, for the expression of CD90 in CD90^+ve^ MuSCs cultures after 3.5 days. DAPI was used to stain the nuclei. Scale bar: 20 μm. A cell doublet with polarized expression of CD90 (dotted area) is magnified in the lower panels. Note the scattered CD90 staining, typical of the organization in lipid rafts (Herrera-Molina et al., 2013). (**G**) Schematic representation of the multiple injury experiment shown in (H) and (I). (**H**) Representative FACS profiles of CD90^+ve^ and CD90^-ve^ MuSCs fractions identified as YFP^+ve^ cells in 1x or 2x injured muscles and uninjured contralateral muscles from tamoxifen-injected *Pax7^CreERT2^;R26R^EYFP^* mice. (**I**) Percentage of CD90^+ve^ MuSCs over total MuSC population in 1x or 2x injured and contralateral muscles, quantified in experiments as in (H). Error bars represent mean ± SEM, n=3.

Firstly, we compared the tendency of CD90^+ve^ and CD90^-ve^ MuSCs to undergo terminal differentiation. Transcripts and protein levels of Mgn were upregulated in CD90^+ve^ compared to CD90^-ve^ MuSCs (**Figures 3B-C**, and **S4A**). These data suggest that the propensity of CD90^+ve^ MuSCs to undergo differentiation (and fusion into the newly regenerated fibers) may be one of the mechanisms restoring the initial 1:1 ratio between CD90^+ve^ and CD90^-ve^ MuSCs after injury resolution.

Furthermore, we conducted experiments to gain insight into their lineage relationship. We sorted CD90^+ve^ and CD90^-ve^ MuSCs from uninjured tamoxifen-injected *Pax7^CreERT2^;R26R^EYFP^*mice and evaluated the expression of CD90 after 3.5 days and 10 days in culture (**Figures 3D**). We regularly reanalyzed the sorted cells to ensure a high degree of purity of the isolated cell populations (**Figures S4B**). Under standard growth conditions (i.e., Ham’s F10 with the addition of 20% FBS and FGF), CD90^+ve^ MuSCs were able to give rise to CD90^-ve^ cells, whereas CD90 was not induced in the progeny of the CD90^-ve^ MuSCs (**Figures 3E**). This suggests that the CD90^+ve^ MuSCs may lie upstream in the lineage hierarchy. In keeping with this observation, we identified doublets of proliferating cells with polarized expression of CD90 in cultures of CD90^+ve^ MuSCs (**Figure 3F** and **S4C**). Further studies will be required to disclose the role of asymmetric division in restoring the ratio between CD90^+ve^ and CD90^-ve^ MuSCs at the end of the regenerative process. Nevertheless, these mechanisms are sufficiently robust to guarantee the renewal of similar proportions of the two subpopulations after multiple injury-regeneration cycles (**Figures 3G-I**). Intriguingly, when the sorted MuSCs subpopulations were grown in a medium that is classically used to grow mesenchymal stem cells and contains lower serum levels (i.e., DMEM with 10% FBS and FGF), the propensity of CD90^+ve^ MuSCs to give rise to CD90^-ve^ cells was blunted (**Figures S4D-E)**. These results indicate that the expression of CD90 during the lineage progression of the CD90^+ve^ MuSCs can be influenced by extrinsic factors associated with the medium, and parallel similar observations obtained in bone marrow or cord blood derived mesenchymal stem cell compartments (Adamzyk et al., 2013; Laitinen et al., 2016). These data suggest that the similarity between CD90^+ve^ MuSCs and mesenchymal stem cells is not limited to the expression of CD90 but also involves certain behavioral aspects, such as the medium preference for *in vitro* maintenance.

### CD90 identifies a subpopulation of human early-activating MuSCs

Next, we evaluated the existence of a subpopulation of MuSCs marked by CD90 in the human muscle. Our FACS analysis confirmed that CD90 marks approximately half of the human MuSCs (**Figures 4A-B**). We confirmed this heterogeneous expression by interrogating published single-cell RNAseq data (Barruet et al., 2020). Unsupervised clustering analysis identified clusters 11 and 15 as being enriched in CD90 expression (**Figures 4C-E**). Both clusters expressed myogenic markers (**Figure 4E**). Intriguingly, the small cluster 15 was the only one characterized by the up-regulation of cell proliferation genes, suggesting that it might be composed of a restricted number of CD90^+ve^ cells that activated during isolation (**Figure 4E**). These findings demonstrate that CD90 identifies a subpopulation of MuSCs in the human muscle. To validate this, we employed a second MuSCs gating strategy that confirmed the separation of the MuSCs pool into two subpopulations based on CD90 expression in homeostatic conditions (**Figure S5A-B**). Importantly, the propensity of CD90^+ve^ MuSCs for rapid activation in human muscle was confirmed by *ex vivo* EdU incorporation (**Figure 4F**). We next applied a muscle injury paradigm and analyzed the tendency of CD90^+ve^ and CD90^-ve^ MuSCs to activate *in vivo* (Jensen et al., 2022) (**Figure 4G**). In agreement with the *ex vivo* data, at early time-points after muscle injury, human CD90^+ve^ MuSCs showed increased activation, measured by augmented cell size and CD82 expression (**Figure 4H** and **S5C**) (Hall et al., 2020). Notably, the proportion of CD90^+ve^ MuSCs significantly rose upon muscle injury, suggesting increased cell proliferation (**Figures 4I-J** and **S5C**).

**Figure 4.**
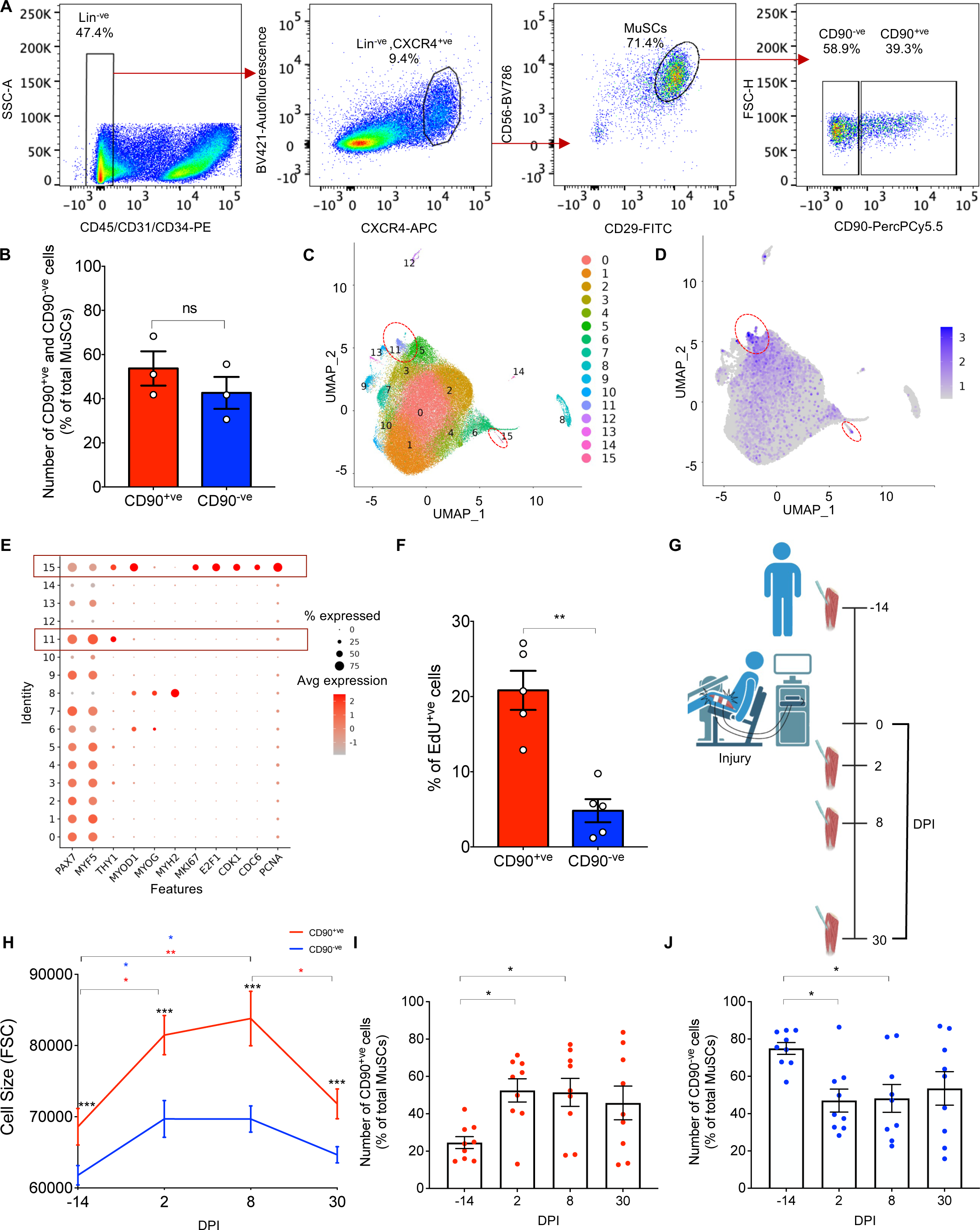
CD90 identifies a subpopulation of human early-activating MuSCs. (**A**) FACS profile of CD90 expression in MuSCs from human muscle negatively selected with CD45, CD31 and CD34 antibodies (referred to as Lin^-ve^), and positively stained with CXCR4, CD29 and CD56 antibodies. (**B**) Quantification of CD90^+ve^ and CD90^-ve^ human MuSCs identified as in (A). Error bars represent mean ± SEM, n=3. (**C**) 2D UMAP plot showing the clusterization of human MuSC subpopulations elaborated from a publicly available scRNA-seq dataset (Barruet et al., 2020). Cells are clustered according to transcriptome similarity. Each dot represents one cell that is colored according to the cluster it belongs. Red-dotted circles indicate CD90-enriched clusters. (**D**) Feature plot showing the localized gene expression of CD90 within the 2D UMAP. CD90-enriched clusters are enclosed in red-dotted circles. (**E**) Dot plot displaying the average expression and percentage of cells expressing myogenic and proliferation markers. Larger dot size indicates more cells expressing each gene, and color indicates the level of expression. Note that the expression of CD90 is enriched in cluster 11 and cluster 15, and proliferation markers are preferentially expressed by cluster 15. (**F**) Percentage EdU^+ve^ human CD90^+ve^ and CD90^-ve^ MuSCs isolated from quadriceps muscle. Samples were fixed 2 days after sorting. Error bars represent mean ± SEM, n=5. (**G**) Schematic representation of human muscle injury model and time-points for collection of muscle biopsies. Muscle injury was induced using electrically stimulated eccentric muscle contractions. (**H**) Cell size (forward scatter) of CD90^+ve^ and CD90^-ve^ human MuSCs before (-14) and 2-, 8- and 30-days post injury (DPI). (**I**, **J**) Relative content of CD90^+ve^ (I) and CD90^-ve^ (J) human MuSCs (% of total MuSCs) before (-14) and 2-, 8- and 30-days post injury (DPI). Error bars represent mean ± SEM, n=9.

### CD90 is a determinant of functional heterogeneity

The expression of CD90 protein is increased in CD90^+ve^ MuSCs during muscle injury (**Figures 5A-B**). To further analyze the functional role of CD90 we silenced CD90 expression in freshly isolated CD90^+ve^ and CD90^-ve^ MuSCs by using a mix of interfering RNAs against CD90 (siCD90) or a scrambled control siRNA (ctrl). Effective CD90 silencing was verified by RT-qPCR and immunostaining (**Figures 5C-E**). CD90 silencing in CD90^+ve^ MuSCs resulted in reduced activation and proliferation, although no significant reduction was detected in the CD90^-ve^ fraction (**Figures 5F-G**). These results demonstrate that CD90 impacts the proliferative potential of a fraction of MuSCs.

**Figure 5.**
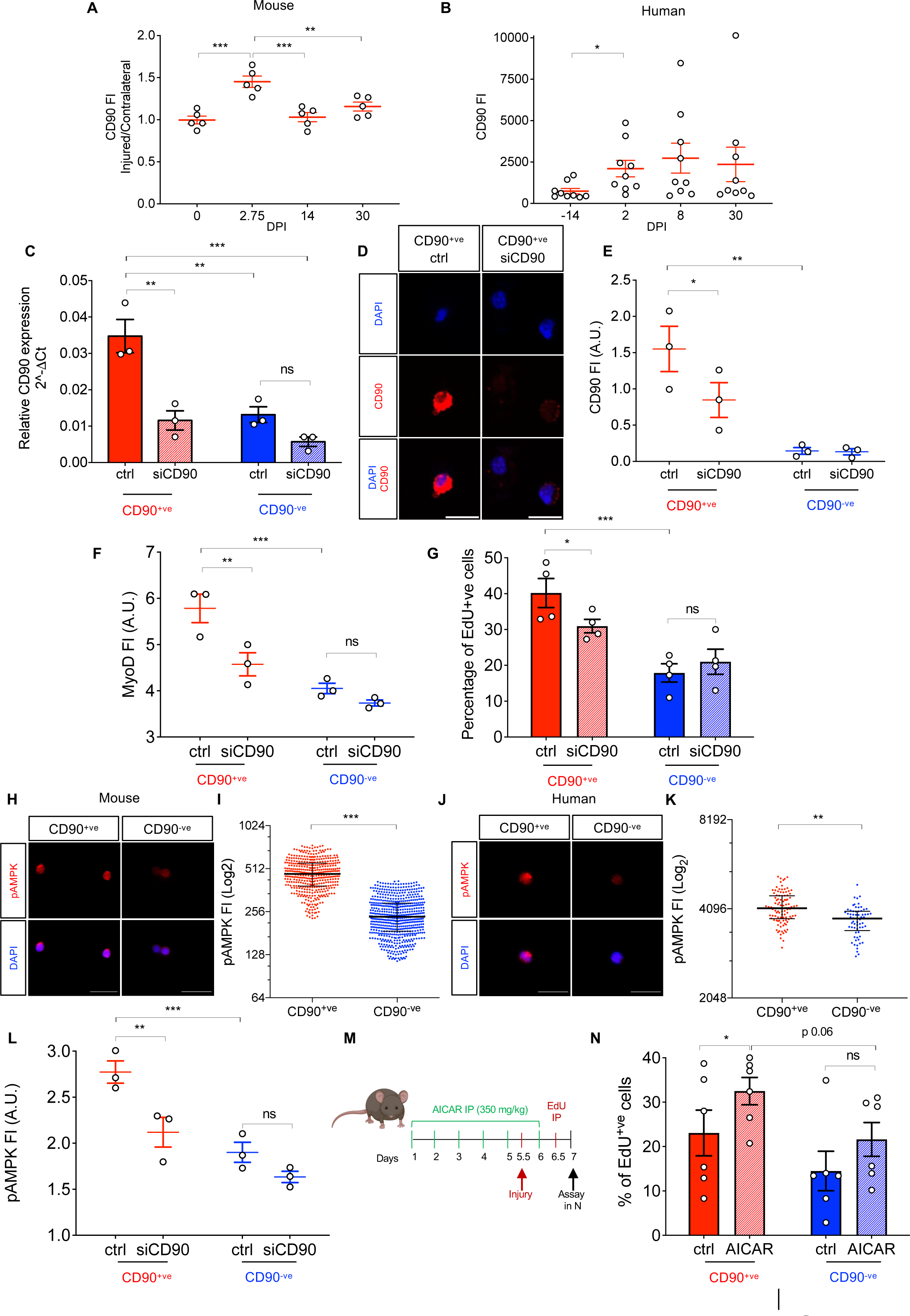
CD90 is a determinant of functional heterogeneity. (**A**, **B**) FACS analysis of CD90 protein expression expressed as median fluorescence intensity (FI) in mouse (A) and human (B) CD90^+ve^ MuSCs at specified time-point of muscle injury (DPI). Error bars represent mean ± SEM, n=5 (A) and 9 (B). (**C**) RT-qPCR analysis of CD90 in CD90^+ve^ and CD90^-ve^ MuSCs treated with CD90-specific siRNA (siCD90) or scrambled control siRNA (ctrl) after 2 days in culture. Error bars represent mean ± SEM, n=3. A two-way Anova test was performed. (**D**, **E**) Images of CD90^+ve^ MuSCs treated with ctrl or siCD90 siRNAs (D), and quantification of CD90 fluorescence intensity (FI) expressed in arbitrary units (A.U.) (E) after 1.5 days in culture. Error bars represent mean ± SEM, n=3. Scale bar: 10 μm (D). A two-way Anova test was performed (E). (**F**, **G**) Immunofluorescence quantification of MyoD fluorescence intensity (FI) (F), and percentage of EdU^+ve^ (G) in FACS isolated CD90^+ve^ and CD90^-ve^ MuSCs treated with ctrl or siCD90 siRNAs after 2.5 days in culture. EdU was administered 2 hours before the end of the treatment. Error bars represent mean ± SEM, n = 3 (F) and 4 (G). A.U.: arbitrary units. A two-way Anova test was performed. (**H**-**K**) Representative images and quantification of ^p^AMPK fluorescence intensity (FI) in CD90^+ve^ and CD90^-ve^ MuSCs freshly isolated from *Pax7^CreERT2/wt^;R26R^EYFP/wt^* mice (H-I) and human muscle biopsies (J-K). Scale bar: 20 μm. In graphs in I and K, each dot represents a cell. Error bars represent the median with interquartile range; the number of cells/replicate is ≥ 400 for mouse and ≥ 60 for human specimens. A two-tailed Mann-Whitney test was performed. See Figure S6A-B for two additional independent experiments. (**L**) Immunofluorescence quantification of ^p^AMPK fluorescence intensity (FI) in FACS isolated CD90^+ve^ and CD90^-ve^ MuSCs treated with ctrl or siCD90 siRNAs after 2.5 days in culture. Error bars represent mean ± SEM, n=3. A.U.: arbitrary units. A two-way Anova test was performed. **(M)** Schematic representation of *in vivo* AICAR treatment. **(N)** Percentage of EdU^+ve^ MuSCs after AICAR administration as described in (M). MuSCs were identified as YFP^+ve^ cells from hindlimb muscles of tamoxifen-injected *Pax7^CreERT2/wt^;R26R^EYFP/wt^* mice. Error bars represent mean ± SEM, n=6.

An increasing body of evidence indicates that the energetic state is pivotal in driving MuSCs’ activation (Relaix et al., 2021). In line with this view, the metabolic sensor AMP-activated protein kinase (AMPK), a master regulator of mitochondrial biogenesis, reportedly plays a key role in modulating MuSCs’ activation and proliferation (Fu et al., 2015; Theret et al., 2017). Notably, CD90 was shown to positively modulate AMPK phosphorylation and consequent activation in ovarian cancer cells (Chen et al., 2016). We, therefore, wondered if AMPK could operate downstream of CD90 also in CD90^+ve^ MuSCs. In agreement with this hypothesis, immunofluorescence experiments demonstrate that both murine and human CD90^+ve^ MuSCs are enriched in the active form of AMPK (**Figures 5H-K and S6A-B**). Importantly, the levels of active AMPK were significantly downregulated upon CD90 silencing in CD90^+ve^ MuSCs (**Figure 5L**). In keeping with the idea that AMPK is operating downstream of CD90 in the control of activation, when we administered the AMPK activator AICAR *in vivo,* an accelerated cell cycle was observed upon muscle injury in MuSCs, which reached the statistical significance for the CD90^+ve^ fraction (**Figures 5M-N)**. A large body of evidence demonstrates that the activation of AMPK signaling and subsequent enhancement of mitochondrial function are key components of the adaptation response to various forms of exercise (Mesquita et al., 2021; Vissing et al., 2013). Intriguingly, when we looked at a cohort of subjects exposed to a protocol of low-load blood flow restricted resistance exercise (BFRRE), which is characterized by modest mechanical stress and increased mitochondrial function, we observed an increase in the amount of CD90^+ve^ MuSCs without changes in the CD90^-ve^ counterpart (**Figures S6C-D)** (Farup et al., 2015; Groennebaek et al., 2018; Wang et al., 2023). Altogether, these observations suggest that CD90 can promote conditions favoring the activation of AMPK and reveal the existence of a CD90-AMPK axis contributing to the propensity for activation shown by CD90^+ve^ MuSCs.

### Collagen VI is preferentially expressed by CD90^+ve^ MuSCs and promotes quiescence

If CD90 has a role in promoting activation and proliferation of CD90^+ve^ MuSCs upon injury, we reasoned that additional CD90-independent mechanisms should be in place to counteract this tendency in the physiological state. We interrogated publicly available human and mouse single-cell RNAseq datasets to gain an insight into this aspect (Barruet et al., 2020; De Micheli et al., 2020). This analysis reveals that the transcriptome of CD90^+ve^ MuSCs is enriched in genes coding for extra-cellular matrix (ECM) components (**Figures S7A** and **S7B**). Of note, Col6 stands out as one of the most differentially expressed ECM components between CD90^+ve^ and CD90^-ve^ MuSCs (**Figure S7C** and **Table S1**). An RT-qPCR analysis for Col6 genes confirmed this differential expression and showed enrichment in quiescent MuSCs (**Figure 6A**). Moreover, immunostaining of *gastrocnemius* muscle sections confirmed differential Col6 protein levels in the niche of CD90^+ve^ and CD90^-ve^ MuSCs (**Figures 6B-C**).

**Figure 6.**
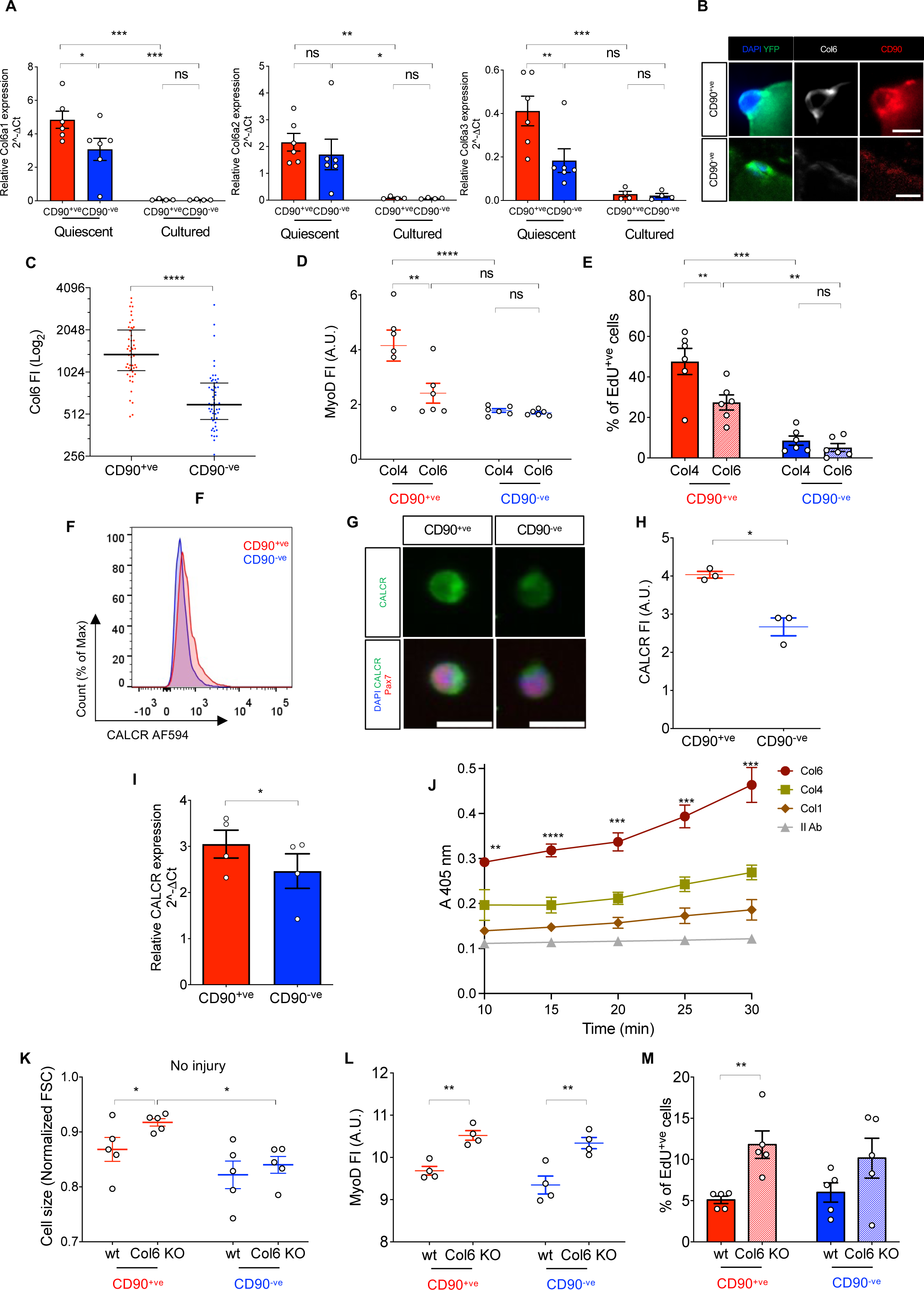
Collagen 6 is preferentially expressed by CD90^+ve^ MuSCs and promotes quiescence. (**A**) RT-qPCR analysis of *Col6a1*, *Col6a2*, and *Col6a3* genes in FACS-purified CD90^+ve^ and CD90^-ve^ MuSCs subpopulations directly isolated (quiescent) from the muscles or cultured for 2 days (cultured). Error bars represent mean ± SEM, n=6 (quiescent) or 4 (cultured) biological replicates. A two-way Anova test was performed. (**B**) Immunofluorescence images of Col6 staining in sections of the *gastrocnemius* muscle of *Pax7^CreERT2/wt^;R26R^EYFP/wt^* mice. Anti-YFP, -CD90 antibodies and DAPI were used to identify CD90^+ve^ (upper panel) and CD90^-ve^ (lower panel) MuSCs and nuclei. Scale bar: 10 μm. (**C**) Quantification of Col6 staining in CD90^+ve^ and CD90^-ve^ MuSCs identified as in (B). Each dot represents a cell. Error bars represent median with interquartile range, the number of cells is ≥ 48. A two-tailed Mann-Whitney test was performed. (**D**, **E**) Quantification of MyoD fluorescence intensity (FI) (D) and percentage of EdU incorporating cells (E) in FACS-isolated CD90^+ve^ and CD90^-ve^ MuSCs after 2.5 days in culture on Col6- or Collagen IV (Col4)-coated dishes. EdU was administered 2 hours before fixation. Error bars represent mean ± SEM, n=6. A.U.: arbitrary units. A two-way Anova test was performed. (**F**) FACS histogram of CALCR protein expression in CD90^+ve^ and CD90^-ve^ MuSCs subpopulations. (**G-H**) Immunofluorescence images of CALCR staining in freshly isolated CD90^+ve^ and CD90^-ve^ MuSCs (Pax7^+ve^) (G) and quantification of CALCR fluorescence intensity (FI) (H). Error bars represent mean ± SEM, n =4. A.U.: arbitrary units. G scale bar: 8 μm. (**I**) RT-qPCR analysis of *CALCR* in FACS-purified CD90^+ve^ and CD90^-ve^ MuSCs subpopulations, identified as YFP^+ve^ cells from hindlimb muscles of uninjured tamoxifen-injected *Pax7^CreERT2/wt^;R26R^EYFP/wt^* mice. Error bars represent mean ± SEM, n=4 biological replicates. (**J**) Enzyme-linked immunosorbent assay (ELISA) showing binding of ColVI and CALCR based on the absorbance of the horseradish peroxidase at specified time-points. P-value refers to the absorbance of ColVI vs. all other conditions. Error bars represent mean ± SEM, n=3. A one-way Anova test was performed. (**K**) Measurement of FSC-A of wild-type (wt) and *Col6a1^–/–^* (Col6 KO) CD90^+ve^ and CD90^-ve^ MuSC cell size, normalized by 15 μm standard beads. Note that the *Col6a1^–/–^* CD90^+ve^ MuSCs display increased cell size compared to their wild-type counterparts. Error bars represent mean ± SEM, n=5. (**L**) Immunofluorescence quantification of MyoD fluorescence intensity (FI) in CD90^+ve^ and CD90^-ve^ MuSCs isolated from wild-type (wt) or *Col6a1^–/–^* (Col6 KO) uninjured hindlimb muscles. Error bars represent mean ± SEM, n =4. A.U.: arbitrary units. (**M**) EdU incorporation in CD90^+ve^ and CD90^-ve^ MuSCs in wild-type (wt) or *Col6a1^–/–^* (Col6 KO) uninjured muscles. EdU was administered 12 before sacrifice. Error bars represent mean ± SEM, n=5.

Recent studies showed that *in vivo* down-regulation of Notch signaling negatively modulates Collagen V and Col6 in muscle, and that genetic ablation of Collagen V induces a break of quiescence in MuSCs, as previously documented for the inhibition of Notch signaling (Baghdadi et al., 2018; Bjornson et al., 2012; Mourikis et al., 2012). Based on these observations, we hypothesized that, similarly to Collagen V, Col6 could promote MuSCs’ quiescence. To investigate this, we first use a gain-of-function approach relying on the over-expression of the Notch intracellular domain (NICD) to confirm that the activation of the Notch signaling promotes the expression of Col6 in myogenic cells (**Figures S7D-E**). Then, we analyzed the kinetics of activation of FACS-purified CD90^+ve^ and CD90^-ve^ MuSCs by using Col6 or Collagen IV as alternative substrates (**Figures 6D-E**). A reduction in MyoD levels and in the number of EdU^+ve^ cells was observed on Col6, compared to Collagen IV (**Figures 6D-E**). This difference is statistically significant in CD90^+ve^ MuSCs, revealing Col6 as a mediator of CD90^+ve^ MuSCs quiescence (**Figures 6D-E**).

Our data suggest that CD90^+ve^ MuSCs exhibit an increased sensitivity to the quiescence-inducing influence of Col6 supplementation (**Figures 6D-E**). To gain insights into this aspect, we looked at the expression of the Calcitonin receptor (CALCR), which is reportedly involved in the control of MuSCs’ quiescence and was recently shown to bind to Collagen V and mediate its quiescence-promoting effects (Baghdadi et al., 2018; Fukada et al., 2007). Flow cytometry and immunofluorescence analysis demonstrate that CALCR is more expressed in CD90^+ve^ MuSCs compared to CD90^-ve^ MuSCs (**Figures 6F-H**). This difference was confirmed at the transcriptional level (**Figure 6I**). Importantly, we also demonstrate that, similarly to Collagen V, Col6 specifically binds to the CALCR (**Figures 6J** and **S7F**).

To further corroborate the role of Col6 in promoting quiescence, we investigated the dynamics of MuSCs activation *in vivo* in Col6 null (*Col6a1^–/–^*) mice. In agreement with previous studies, Col6 ablation led to MuSC activation (Urciuolo et al., 2013). Although both CD90^+ve^ and CD90^-ve^ MuSCs appeared to activate in response to Col6 ablation, specific activation parameters, such as cell size, only significantly increased in CD90^+ve^ MuSCs (**Figure 6K-M**). The robust response of CD90^+ve^ MuSCs may be explained by the intrinsic propensity of CD90^+ve^ MuSCs to activate (see above). To validate the quiescence-promoting role of Col6, we investigated the fate of activated MuSCs in *Col6a1^–/–^* mice by marking them *in vivo* with multiple injections of EdU. Histological analysis of serial muscle sections revealed that EdU^+ve^ nuclei were readily identifiable in newly formed myofibers (eMyHC^+ve^) and in differentiating MyoD^+ve^/Mgn^+ve^ cells. This suggests that activated MuSCs progress toward terminal differentiation in *Col6a1^–/–^* mice (**Figure S8A-B**).

Altogether these findings underline an important role exerted by Col6 in MuSCs during quiescence and suggest that its loss may favor MuSCs transition from quiescence to differentiation. This phenomenon phenotypically resembles what occurs when the blocking of Notch and Collagen V signaling breaks MuSCs’ quiescence (Baghdadi et al., 2018; Bjornson et al., 2012; Mourikis et al., 2012). The preferential expression of Col6 and its binding partner CALCR in CD90^+ve^ MuSCs, extends this concept in the context of MuSCs’ heterogeneity and indicates that distinct MuSCs subpopulations may propagate different cell-autonomous signaling in the local niche to ensure quiescence.

### The fraction of CD90^+ve^ MuSCs is reduced in muscular dystrophy

The idea that Col6 may differentially affect quiescence and lineage progression of CD90^+ve^ and CD90^-ve^ MuSCs is corroborated by the finding that *Col6a1^–/–^* mice exhibited a perturbed proportion between the two subpopulations (**Figures 7A-B**). In *Col6a1^–/–^*mice, the number of CD90^+ve^ MuSCs was significantly lower compared to the CD90^-ve^ counterpart (**Figure 7C**).

**Figure 7.**
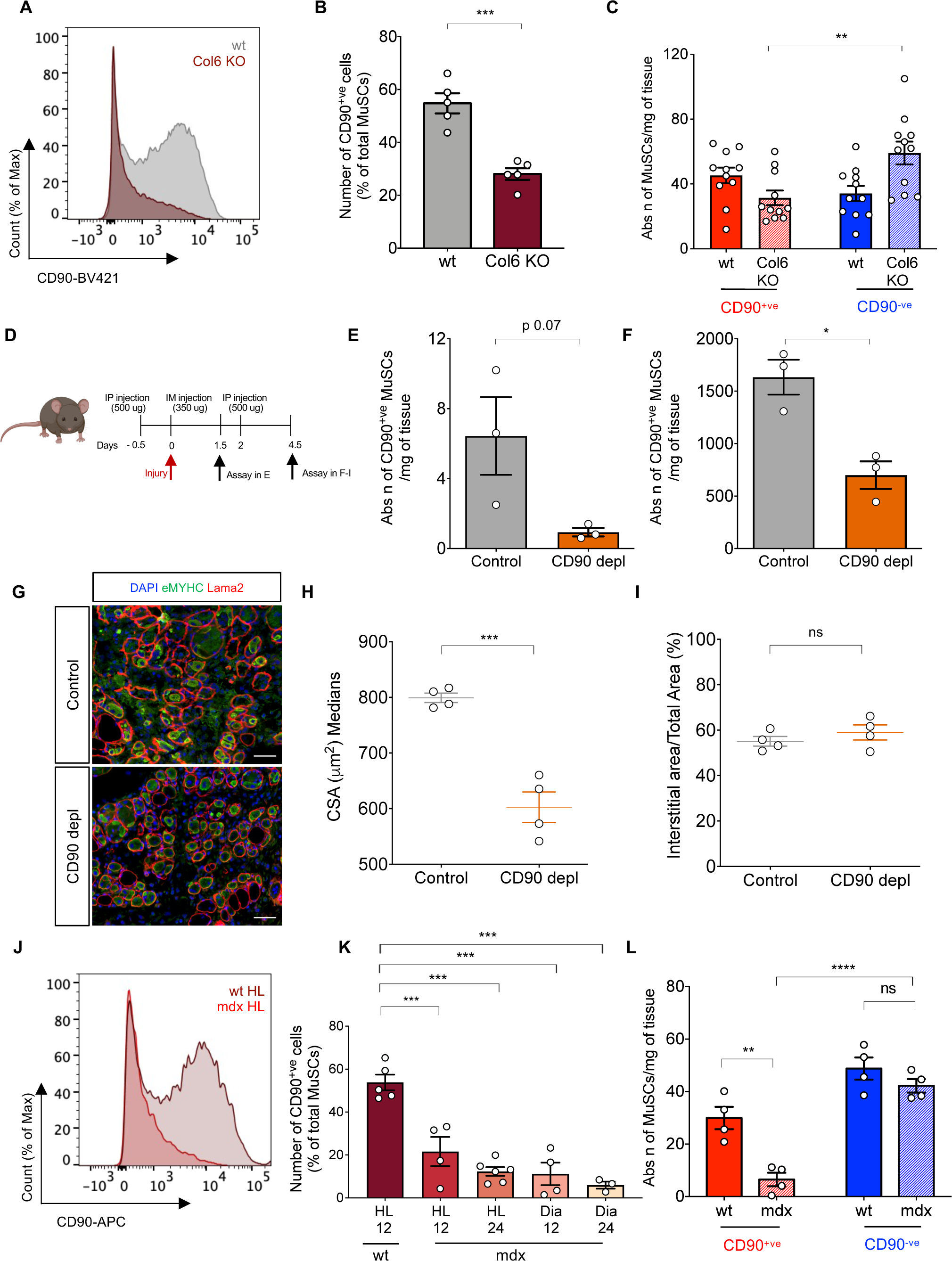
The fraction of CD90^+ve^ MuSCs is reduced in muscular dystrophy. (**A**) FACS histograms showing the distribution of CD90 staining in MuSCs of hindlimb muscles from 6 months-old wild-type (wt) and *Col6a1^–/–^* (Col6 KO) mice. *Col6a1^–/–^* MuSCs display reduced expression of CD90. (**B**, **C**) Quantification of CD90^+ve^ cells expressed as a percentage of the total MuSC population (B), and absolute numbers of CD90^+ve^ and CD90^-ve^ MuSCs per mg of tissue (C) in hindlimb muscles of 6 months-old wild-type (wt) and *Col6a1^–/–^* (Col6 KO) mice. Error bars represent mean ± SEM, n =5 (B) and n=11 (C). A two-way Anova test performed in (C). (**D**) Schematic representation of CD90^+ve^ cells depletion. (**E, F**) Absolute numbers of CD90^+ve^ per mg of tissue in hindlimb muscles of mice treated with isotype control antibodies or antibodies depleting CD90^+ve^ cells at 1.5 days post-injury (E) and 4.5 days post-injury (F). (**G**) Immunostaining with embryonic myosin heavy chain (eMyHC)- and laminin-recognizing antibodies of *gastrocnemius* muscles of mice treated with isotype control antibodies and antibodies depleting CD90^+ve^ cells at 4.5 days post-injury. DAPI was used to stain the nuclei. Scale bar: 50 μm. (**H, I**) Cross-sectional area (CSA) of the eMyHC^+ve^ fibers (H) and interstitial area (I) quantifications of the muscles as in (G). Error bars represent mean ± SEM, n =4 biological replicates. (**J**) FACS histograms showing the distribution of CD90 in MuSCs from 12-month-old wild-type (wt) and *mdx* mice. (**K**) Proportion of CD90^+ve^ cells over total YFP^+ve^ MuSCs in hindlimb (HL) and diaphragm (Dia) muscles of *Pax7^CreER/wt^;R26R^YFP/wt^* 12-month-old wild-type (wt) mice, and of 12- or 24-months old *mdx* mice. Error bars represent mean ± SEM, n ≥ 3. A one-way Anova test was performed. (**L**) Quantification of CD90^+ve^ and CD90^-ve^ MuSCs in hindlimb muscles of wild-type (wt) and *mdx* mice identified as in (K) and expressed as the absolute number of cells per mg of tissue. Error bars represent mean ± SEM, n =4. A two-way Anova test was performed.

Mutations of Col6 genes in humans are causative for a distinct group of inherited muscle disorders, whose major forms are Bethlem myopathy and Ullrich congenital muscular dystrophy (Lampe and Bushby, 2005). Furthermore, *Col6a1^–/–^*mice are characterized by defective muscle regeneration and display signs of muscular myopathy (Urciuolo et al., 2013). To investigate the possibility that the imbalance between CD90^+ve^ and CD90^-ve^ MuSCs may contribute to impaired muscle regeneration, we administered *in vivo* antibodies, previously used by others to deplete CD90^+ve^ cells in other tissues, and evaluated the muscle repair process (**Figure 7D**) (Powell et al., 2012; Zhou et al., 2022). FACS analysis demonstrated effective depletion of CD90^+ve^ cells in muscles at different time-points after injury (**Figures 7E-F**). Notably, a delay in regeneration was noticeable in depleted animals, as demonstrated by a reduced size of newly regenerated fibers without significant changes in the interstitial portion of the muscle at 4.5 days after injury (**Figures 7G-I**).

Our findings prompted us to investigate if the balance between CD90^+ve^ and CD90^-ve^ MuSCs is also altered in other inherited muscle pathologies characterized by defective regeneration. We observed a marked alteration in the ratio between the two subpopulations in the *mdx* mouse model of Duchenne muscular dystrophy (**Figures 7J-L**). Indeed, *mdx* mice displayed a specific loss of CD90^+ve^ MuSCs, which was exacerbated by aging (**Figure 7K**), in line with the progressive features of the dystrophic phenotype (Pastoret and Sebille, 1995). Moreover, the disappearance of the CD90^+ve^ MuSCs was dramatic in the diaphragm muscle (**Figure 7K**), which is characterized by a severe dystrophic pathology (Stedman et al., 1991). Altogether, these data support a link between impaired regeneration and loss of the physiological balance between CD90^+ve^ and CD90^-ve^ MuSCs in dystrophic muscles.

## DISCUSSION

The transition from quiescence to proliferation in stem cells needs to be controlled carefully, as misregulation can lead to defects in cell growth, tissue homeostasis, or regeneration. Quiescent stem cells are poised in an inactive state in anticipation of proliferation and differentiation (van Velthoven and Rando, 2019). Nevertheless, the signals controlling the delicate balance between quiescence and activation remain poorly understood. Although recent studies have highlighted the heterogeneity of stem cells in various tissues, information about how quiescence is interpreted in this context is lacking. Our current data demonstrate that quiescence is not homogeneously controlled by the heterogeneous pool of stem cells resident within the skeletal muscle. Instead, subpopulation-specific mechanisms govern the balance between quiescence and activation in both mouse and human MuSCs. Our data reveal the existence of two signaling axes governing two distinct but complementary aspects of CD90^+ve^ MuSCs. On the one hand, our results show that CD90 expression grants a fraction of MuSCs a distinct molecular program involving the metabolic sensor AMPK and ensuring different kinetics of recruitment upon injury. Our results indicate that an active CD90-AMPK axis grants CD90^+ve^ MuSCs a higher propensity for activation upon injury. This calls for future mechanistic studies to comprehend the molecular link between CD90 and AMPK. On the other hand, we demonstrate here that CD90^+ve^ MuSCs can produce relatively high levels of Col6 and propagate their autonomous cell signaling in the local niche to ensure quiescence in the absence of muscle damage. We documented that Col6 binds to the CALCR, which is also upregulated in CD90^+ve^ MuSCs, and propose that Col6 mediates its quiescence-promoting effects through this interaction. To our knowledge, this study is the first to demonstrate that the propensity for activation and the need for quiescence maintenance must be orchestrated and are differentially interpreted in distinct subpopulations of stem cells.

A number of studies have started to elucidate how the niche ECM modulates MuSCs’ lineage progression by interacting with receptors on the MuSCs to control cell cycle entry and cell division (Baghdadi et al., 2018; Bentzinger et al., 2013; Urciuolo et al., 2013). Our finding that Col6 and CALCR are preferentially expressed by CD90^+ve^ MuSCs expand our previous study documenting Col6 production by quiescent MuSCs (Urciuolo et al., 2013). Notably, the impact of Col6 on the transition from quiescence to activation appears to be unequal between CD90^+ve^ and CD90^-ve^ MuSCs (i.e., activation parameters in the presence of Col6, cell size increase, and cell number changes in *Col6a1^–/–^*). This is possibly due to the enhanced expression of CALCR in CD90^+ve^ MuSCs, but we cannot exclude the contribution of other unknown mechanisms. In this regard, it should be considered that Col6 and CALCR are not entirely absent in quiescent CD90^-ve^ MuSCs (see **Figure 6A** and **G**), and Col6 is reportedly also expressed by other muscle-resident cells (Castagnaro et al., 2018). Therefore, it is not surprising that the modulation of Col6 is also impacting the dynamics of this subpopulation to a certain extent. Altogether, our data support the idea that the niche may be heterogeneously formed and deciphered by individual MuSCs, thereby contributing to their own phenotypic diversification.

Several lines of evidence suggest that MuSCs become exhausted and lose their regenerative potential with the progression of the dystrophic disease, thereby contributing to the severity of this condition (Biressi and Gopinath, 2015). Studies carried out by different groups, including ours, indicate that MuSCs present abnormalities in dystrophic muscle (Biressi et al., 2014; Delaporte et al., 1990; Jasmin et al., 1984; Pessina et al., 2015; Rhoads et al., 2013). The molecular mechanisms underlining these changes are poorly explored, and it needs to be clarified to what extent they reflect alterations in the muscle environment or intrinsic changes in the MuSCs. Importantly, dystrophin, the product of the gene mutated in Duchenne muscular dystrophy, is present in both fiber sarcolemma and MuSCs, where it controls asymmetric division and MuSCs’ lineage decisions (Dumont et al., 2015). Our data show that CD90 is asymmetrically segregated in proliferating MuSCs, that CD90^+ve^ MuSCs can give rise to their CD90^-ve^ counterparts, and that the balance between CD90^+ve^ and CD90^-ve^ MuSCs is perturbed in dystrophic muscles. These findings support a contention that the severity of the dystrophic condition is influenced by MuSC alterations (Ganassi et al., 2021).

Within muscle, CD90 heterogeneity is not unique to MuSCs. CD90 was described in stromal cells with myogenic capacity (Crisan et al., 2008). High levels of CD90 expression were shown to mark a subpopulation of non-myogenic fibroblast-like cells in adult muscle (Fukada et al., 2008). We show that CD90 is heterogeneously marking MuSCs and FAPs within murine and human muscle. Notably, a CD90^+ve^ subpopulation of FAPs predominates within the muscles of diabetic patients (Farup et al., 2021). Similarly to CD90^+ve^ MuSCs, CD90^+ve^ FAPs appear to express ECM components more robustly and enter the cell cycle more rapidly than CD90^-ve^ FAPs (Farup et al., 2021). Our data disclose that CD90 is not only a marker but also a molecular determinant of MuSCs phenotypic diversification and call for further investigation of its functional role in FAPs. The expression of CD90 in multiple cell types suggests that CD90 might play a role in orchestrating the cellular responses involved in tissue repair.

By understanding the molecular events controlling MuSCs’ quiescence, our data may provide a step forward in comprehending the mechanisms controlling quiescence in other stem cell compartments, particularly in the context of heterogeneity. Notably, CD90 is expressed in the long-term repopulating fraction of hematopoietic progenitors (Szilvassy et al., 1989). Future studies will ascertain to what extent mechanisms controlling heterogenic stem cell lineage decisions are conserved among tissues.

## MATERIALS AND METHODS

### Animal models and procedures

Mice were purchased from Jackson labs through Charles River. Unless otherwise stated, mice in the *C57BL/6J* background (stock no. 000664) were used as wild-type (wt) animals. *Pax7^CreERT2^* (stock no. 017763) and *Myf5^CreER^* (stock no. 023342) homozygous males were crossed with *R26R^EYFP/EYFP^* females (stock no. 006148) to obtain respectively *Pax7^CreERT2/wt^*;*R26^EYFP/wt^* and *Myf5^CreER/wt^*;*R26^EYFP/wt^* experimental animals in the *C57BL/6J* background. For specific experiments, *Myf5^CreER^*;*R26^EYFP^*mice were bred to *FVB* mice (stock no. 001800) for at least 3 generations to obtain experimental animals in the *FVB* background. To obtain *Pax7^CreERT2/wt^*;*R26^LacZ/wt^*experimental animals, *R26R^LacZ/LacZ^* females (stock no. 003474) were crossed with *Pax7^CreERT2^* homozygous males. *B6Ros.Cg-Dmd^mdx–4Cv/J^* (herein referred to as *mdx*; stock no. 002378) homozygous female mice were crossed with *Pax7^CreERT2/wt^;R26R^EYFP/wt^*males to obtain *Pax7^CreERT2/wt^;R26R^YFP/wt^;mdx* male experimental mice. *Col6a1*^–/–^ mice in the *C57BL/6J* background were previously described (Urciuolo et al., 2013).

Tamoxifen (MP Biomedicals, 50 mg/ml, 100 μl/injection) was administrated intraperitoneally twice a week for 2 weeks. Unless otherwise stated, for *in vivo* proliferation assays, 100 μl of 15 mM EdU (Roth) were injected intraperitoneally 12 hours before sacrifice. For *in vivo* experiments with *Col6a1*^–/–^ mice, EdU was injected for 5 consecutive days before analysis. AICAR (5-aminoimidazole-4-carboxamide ribonucleotide, Toronto Research Chemicals) was administered intraperitoneally at 350 mg/kg body weight for 6 consecutive days, with the last injection being performed 12 hours after muscle injury.

Muscle injury was performed by repeated punctures (approximately 100) with 29-gauge needle on lower limb muscles to cover the whole muscle surface. Animal care and experimental procedures were conducted in accordance with the Ethical Committee of the University of Trento and were approved by the Italian Ministry of Health (Authorization No. 228/2017-PR).

### Murine MuSCs isolation and staining

Isolation of MuSCs by enzymatic dissociation was performed as previously described (Florio et al., 2022). For isolation of MuSCs from *Col6a1*^–/–^ mice, muscles were stored for 3 hours at 4°C before digestion.

MuSCs were isolated using a FACS Aria II or III (BD) as YFP^+ve^ cells from *Pax7^CreERT2/wt^*;*R26R^EYFP/wt^* mice in the *C57BL/6J* or *mdx* background or purified by positive selection with an anti-VACAM antibody and negative selection with antibodies recognizing CD31, CD45, and Sca1 (Lin^−ve^), as previously described. FAPs as CD45^-ve^/CD31^-ve/^Sca1^+ve^ cells (Florio et al., 2022). Unless otherwise stated, the clone 53-2.1 anti-CD90.2 antibody was used. A list of FACS antibodies is enclosed in **Table S2**. Sorted CD90^+ve^ and CD90^-ve^ cells were routinely analyzed by flow cytometry immediately after sorting to ensure high sorting purity (i.e., > 95%).

For detection of CALCR, the muscle single cell suspension was fixed with 1% paraformaldehyde (PFA) at room temperature (RT) for 15 minutes. Cells were immersed in permeabilization buffer (1% saponin, 0.5% BSA) and stained with conjugated anti-CD90 and not-conjugated anti-CALCR antibodies in cell staining buffer (0.5% BSA) for 40 minutes at 4°C. Samples were incubated with conjugated secondary antibody for 30 minutes at 4°C.

For mitochondrial evaluation, MitoTracker Deep Red (Invitrogen) was added to muscle digests at the concentration of 20 nM and incubated for 15 minutes at 37 °C.

### MuSCs flow cytometry analysis from fixed tissue

Isolation of MuSCs from *in situ* fixed tissue was performed as previously described with minor modifications (Machado et al., 2018). Briefly, hindlimb muscles of *Pax7^CreERT2/wt^*;*R26R^EYFP/wt^* mice were finely minced immediately after dissection in ice-cold 0.5% PFA, and placed in ice-cold 0.5% PFA on a rotating wheel at 4°C for 1 hour. The muscle suspension was then washed and digested in 500 U/ml Collagenase II at 37°C for 1 hour. The muscle suspension was then incubated with 0.8 μg/ml Proteinase K (ThermoFisher) at 37°C for 20 minutes. The muscle suspension was passed through 18 and 19-gauge needles and filtered through 70 µm and 40 µm cell strainers.

### Human muscle biopsies

Human muscle biopsies employed in this study were obtained from adult individuals undergoing surgery in the lumbar region at Ca’ Granda Ospedale Maggiore Policlinico in Milan, which participate to the Telethon Network of Genetic Biobanks (Protocol No. 1504), or donors undergoing leg amputation at Aarhus University Hospital, Denmark. Amputations were due to peripheral leg ischemia (biopsies were obtained from non-ischemic areas and used for *ex vivo* EdU incorporation). The protocol was approved by the local ethical committee of Central Denmark Region (1-10-72-253-16). Procedures were performed under the International Conference on Harmonization of Good Clinical Practice guidelines, the Declaration of Helsinki (2008), and the European Directive 2001/20/EC. Informed consent was obtained after the nature and possible consequence of the study were explained.

The human muscle injury study was an investigator-initiated randomized, double-blinded, placebo-controlled trial with a primary endpoint reported elsewhere (Jensen et al., 2022). The study was conducted in accordance with the Declaration of Helsinki and national laws after approval by the local Research Ethics Committee in Central Denmark Region (1-10-72-301-18) and registered at clinicaltrials.gov (NCT03754842). Participants received oral and written information before written consent was obtained. Skeletal muscle biopsies were obtained from the *vastus lateralis* muscle by a Bergström needle under sterile conditions. Baseline biopsies were obtained before supplementation with Nicotinamide Riboside+Pterostilbene or placebo. Only baseline biopsies (n=26) or complete placebo group biopsies post injury (n=9) were included in present analysis.

The human exercise study was a randomized controlled trial with a primary endpoint reported elsewhere (Wang et al., 2023). The study was undertaken in accordance with the Declaration of Helsinki and approved by the Central Denmark Region Committees on Health Research Ethics (1-10-72-169-20) and registered at ClinicalTrials.gov (NCT04712955). Participants received oral and written information before written consent was obtained. Skeletal muscle biopsies were obtained from the *vastus lateralis* muscle. Details about the subjects are indicated in **Table S3.**

### Human MuSCs isolation and staining

Human MuSCs were isolated by using two strategies. In the first strategy, human muscle tissue was stored overnight at 4°C in DMEM, minced and incubated in Collagenase II at 500 U/ml at 37°C for 1 hour in a shaking water bath. The muscle digest was washed, resuspended in 2000 U/ml Collagenase II and 11 U/ml Dispase, and incubated in a shaking water bath at 37°C for 30 minutes. The muscle suspension was passed through 18 and 19-gauge needles, washed and filtered through 40 μm cell strainer. MuSCs were identified as negative for the expression of CD31/CD34/CD45, and positive for CXCR4/CD29/CD56 (Barruet et al., 2020). In the second strategy, muscle biopsies were transported to the laboratory in ice-cold Hams F10. Percutaneous biopsies collected following muscle injury was placed directly in C-tubes (Miltenyi Biotec) without further manipulation, whereas open biopsies were dissected free of visible tendon/connective tissue and fat and minced. The muscle slurry was transferred to C-tubes containing 8ml wash-buffer. From this point, needle and open biopsies were treated similarly. 700 U/ml Collagenase II and 3.27 U/ml Dispase II were added to the C-tubes, and mechanical and enzymatic muscle digestion was performed at 37°C on the gentleMACS with heaters (Miltenyi Biotec) using a 60 min skeletal muscle digestion program (37C_mr_SMDK1). The suspension was washed and filtered through a 70 μm cell strainer. The cell pellet was resuspended in freezing buffer (StemMACS, Miltenyi Biotec) and stored at -80°C. The cell suspension was thawed on ice 1.5h before FACS. MuSCs were identified as negative for CD31/CD34/CD45, and positive for CD82/CD56 (Billeskov et al., 2023; Farup et al., 2021; Jensen et al., 2021). Human-specific anti-CD90 antibodies were used to identify CD90^+ve^ or CD90^-ve^ MuSCs (**Table S2**). MuSCs were isolated using a FACS Aria III (BD).

### MuSCs *in vitro* analysis

CD90^+ve^ and CD90^-ve^ MuSCs were sorted separately in HAM’s F10 medium supplemented with 20% FBS. Unless otherwise stated, cells were plated into eight-well chambers (Millicell) or 96- or 384-well plates with optical bottom (PerkinElmer) pretreated with 0.1 mg/ml poly-D-lysine (Millipore), followed by coating with 0.1 mg/ml ECM (Sigma). For lineage experiments, 10,000 freshly isolated CD90^+ve^ and CD90^-ve^ MuSCs were seeded into 8-well chambers. For proliferation assay on Col6 or Collagen IV, 10,000 freshly isolated CD90^+ve^ and CD90^-ve^ MuSCs were plated on 96-well plates coated with 2.5 μg/ml Laminin (Invitrogen) and 0.1 mg/ml Col6 or Collagen IV (CliniScience). For activation/proliferation experiments, cells were cultured in HAM’s F10 medium supplemented with 20% FBS and 2.5 ng/ml recombinant human FGF (PeproTech). When indicated, DMEM medium supplemented with 10% FBS, and 2.5 ng/ml recombinant human FGF (PeproTech) was used. For *in vitro* EdU incorporation assay, murine MuSCs were plated for 2.5 days and cultured with medium containing 10 μM EdU (Roth) 2 hours before fixing. For EdU incorporation, human MuSCs were plated for 2 days and 10 μM EdU was immediately added after sorting. EdU incorporation was visualized by Click-iT (Invitrogen) according to the manufacturer’s instructions. For coculture experiments, CD90^+ve^ and CD90^-ve^ MuSCs were isolated from uninjured wt and *Pax7^CreERT2^/R26^REYFP^*mice and plated in 384-well tissue culture plates. The experimental groups included 5,000 CD90^+ve^ wt cells, 5,000 CD90^-ve^ wt cells, 2,500 CD90^+ve^ wt cells cocultured with 2,500 YFP^+ve^ CD90^-ve^ cells, and 2,500 CD90^-ve^ wt cells cocultured with 2,500 YFP^+ve^ CD90^+ve^ cells. For the activation and proliferation analysis, only the wt cells were considered, using the YFP^+ve^ cells to distinguish between the different cell populations in the coculture.

### RT-qPCR

RNA was extracted using TRIzol (Invitrogen) and reverse transcribed using High-Capacity cDNA Reverse Transcription kit (Applied Biosystems) according to the manufacturer’s instructions. Quantitative RT-PCR was performed on a CFX96 Touch^TM^ thermocycler (Biorad) using PowerUp^TM^ SYBR Green Master mix (Invitrogen). Relative quantification was normalized to mouse HPRT (hypoxanthine-guanine phosphoribosyltransferase). A list of primers used is enclosed in **Table S4**.

### Single fiber experiments and whole muscle β-galactosidase staining

Single fibers were isolated from *soleus* muscles and fixed immediately, as described (Biressi et al., 2013). Fibers were fixed in 2% paraformaldehyde for immunofluorescence analysis with anti-YFP and CD90-antibodies. β-galactosidase analysis was performed as previously described (Biressi et al., 2013). Images were acquired using Axio Imager M2 (Zeiss).

### Histology, H&E staining, immunofluorescence and quantitative microscopy

Unless otherwise stated, cells, single fibers and muscle sections were processed for hematoxylin/eosin (H&E) staining and immunofluorescence with standard protocols. A list of primary and secondary antibodies is enclosed in **Table S5** and **Table S6,** respectively. For CD90 staining on muscle sections, muscles were fixed for 4 hours using 0.5% electron microscopy-grade PFA, transferred to 30% sucrose overnight, and frozen in optimum cutting temperature compound (OCT) (Histo-line Laboratories). For immunofluorescence staining of *Col6a1*^–/–^ mice, muscles were flash frozen in isopentane. Muscles were cryosectioned at 6 µm. For the evaluation of the intensity of the CD90 staining, cells were fixed with 4% electron microscopy-grade PFA for 7 minutes at 4°C and incubated with CD90.2-APC antibody or APC-conjugated isotype antibody at RT for 1 hour or at 4°C overnight.

Immunofluorescence imaging was performed using Zeiss Axio Observer (Carl Zeiss), TCS SP8 confocal microscope (Leica), Operetta High Content Imaging System (PerkinElmer), or ImageXpress High Content Confocal Imaging System (Molecular Devices). In the distribution of cells, the extreme 2% of the population was excluded. The mean pixel intensity, expressed as fluorescence intensity (FI), was calculated for each cell using Fiji/ImageJ software (NIH), Harmony 4.1 software (PerkinElmer) or MetaXpress software (Molecular Devices). To test the positivity of the cells for CD90 in lineage experiments, an analysis sequence was set up using Harmony 4.1 software (PerkinElmer). Briefly, the background intensity of areas devoid of cells was subtracted and the CD90 isotype control antibody was used to set a threshold to determine the positivity for CD90 on single cells.

### RNA interference

For small-interfering RNA (siRNA)-mediated CD90 knockdown, 10,000 freshly isolated CD90^+ve^ and CD90^-ve^ MuSCs were plated on 384-well plates coated with 2.5 μg/ml Laminin (Invitrogen) and 0.1 mg/ml Col6 in antibiotic-free media. After plating, CD90^+ve^ and CD90^-ve^ MuSCs were transfected with scrambled siRNA or siRNA targeting CD90 (Dharmacon) at the final concentration of 125 mM using DharmaFECT transfection reagent (Dharmacon). CD90-targeting siRNAs are indicated in **Table S7**. CD90 expression was measured by immunofluorescence 1.5 days post-transfection, and MyoD expression and EdU incorporation was performed 2.5 days post-transfection as previously described.

### Enzyme-Linked Immunosorbent Assay (ELISA)

An indirect ELISA assay was performed using purified CALCR (Abbexa) on plates coated with collagens. Briefly, 96-well Thermo Fisher NUNC MaxiSorp Immuno-plates were coated with 0.1 mg/ml of either ColVI, ColIV, or ColI overnight at 4°C. For CALCR titration, the wells were incubated with 0.001, 0.01, or 0.1 µg of CALCR. In the binding assay shown in Fig. 6J, the wells were incubated with 0.5 µg of CALCR. After incubation, wells were incubated with a primary antibody against CALCR (BioRad cat. AHP3115, 1:2000) for 1 hour at RT. Wells were treated with an anti-alkaline phosphatase antibody (Sigma cat. A3687, 1:2000) and DEA buffer solution (1M diethanolamine, 0.5 mM MgCl_2_, pH 9.8) containing 3 mg/ml of alkaline phosphatase substrate (Sigma). Absorbance was measured at 405 nm (Varioskan LUX reader).

### *In vivo* CD90 depletion

For in vivo CD90 depletion, mice were injected with 500 μg anti-CD90.2 (30H12, BioXCell or BioLegend) or isotype (BioLegend) antibodies IP before needle injury, followed by an intramuscular injection of 350 μg after injury. For CSA and fibrotic area measurements at 4.5 days post-injury, an additional 500 μg IP injection was administered 2.5 days post-injury.

### Cloning and cell transfection

Notch intracellular domain (NICD) was amplified by PCR from pcDNA3 plasmid containing intra-cellular Notch1 (courtesy of G. Consalez) using primers incorporating KpnI and AgeI restriction enzyme sites. The PCR product was cloned into the pmiRFP670-N1 vector (Addgene #79987) using the same restriction enzymes. Transfection was performed with Lipofectamine 3000 (Invitrogen) according to the manufacturer’s instructions, and transfected cells were analyzed 48 hours after plating.

### Analysis of CyTOF and single-cell RNA sequencing

CyTOF data were downloaded from Flowrepository.org (accession code FR-FCM-ZY3E)(Porpiglia et al., 2017). Cell type annotation, gating and tSNE dimensional reduction were performed using the OMIQ software from Dotmatics (www.omiq.ai, www.dotmatics.com). TSNE plot_was generated using the following parameters and markers. Parameters: 1000 Iterations, 250 early exaggeration Iters,5000 learning rate, 30 Perplexity, 0.5 theta, 9754 as seed. Markers: CD34, CD44, Th1.2, Alpha7 Integrin, Mac-1, CD45, VCAM-1, Sca1, Pax7, CD98, CD31. Populations were annotated based on the following markers: Endothelial CD45^-ve^ CD31^+ve^; T Cells CD45^+ve^ CD11b^-ve^ CD90^+ve^; B cells CD45^+ve^ CD11b^-ve^ CD90^-ve^; Macrophages CD45^+ve^ CD11b^+ve^ CD90^-ve^; FAPS CD45^-ve^ CD31^-ve^ ITGA7^-ve^ SCA1^+ve^, Tenocytes CD45^-ve^ CD31^-ve^ ITGA7^-ve^ SCA1^-ve^, MuSCs CD45^-ve^ CD31^-ve^ ITGA7^+ve^ SCA1^-ve^,VCAM1^+ve^,SMMCS CD45^-ve^ CD31^-ve^ ITGA7^-ve^ SCA1^-ve^ VCAM1^-ve^.

Single cell-RNA sequencing (scRNA-seq) analysis was carried out using Seurat suite version 3.0 in R studio (Stuart et al., 2019). Briefly, the mouse dataset was obtained from SRA using SRAtoolkit (Leinonen et al., 2011)(GSE143437). The data was filtered to for cells expressing >200 genes/cell and >20 cell/gene. Integration of the different runs was performed via SCTtransform as previously described (Stuart et al., 2019). ALRA was used through the Seurat wrapper with default settings to obtain the imputed values on the integrated values (Linderman et al., 2022). The correlation coefficient was calculated with reference to CD90. Correlation coefficient was for each gene was plotted using ggplot (Wickham, 2009). scRNA-seq data on human dataset have been integrated and analyzed as described by Barruet and colleagues (Additional file, Source code 1 https://cdn.elifesciences.org/articles/51576/elife-51576-code1-v2.r, (Barruet et al., 2020)). Briefly, data was filtered for cells expressing >500 and <6000 genes and <10% of mitochondrial genes. Integration of the different datasets was performed using FindIntegrationAnchors() and IntegrateData() using the first 30 dimensions. For downstream analysis the integrated dataset was scaled regressing for “nCount_RNA” and “percent.mt” variables, UMAP embedding and neighbors’ identification were performed using the first 30 dimensions. Clustering analysis was performed with a resolution of 0.5. Differential gene expression for CD90 enriched MuSCs cluster (cluster 11) was performed using the FindMarkers() function with the following parameters (min.pct = 0.25, only.pos = FALSE, test.use = “MAST”). Gene Ontology and Reactome analysis were performed on Enricher Platform^57^.

### Statistical analysis and figure creation

Unless stated otherwise, significance was calculated using two-tailed paired Student’s *t*-tests. Statistical analyses were performed using Microsoft Excel or GraphPad Prism softwares. Relevant p-values are indicated in the graphs as follows: *, p≤0.05; ** p≤0.01; *** p≤0.001; ns ≥0.15. Analysis of flow cytometry data was performed using FlowJo software. Scientific drawings were created with Biorender.

## ACKNOWLEDGMENTS

We thank Benoit Viollet, Toma Tebaldi, Giorgia Bucciarelli, Giulio Cossu, Alessandro Magli and Luciano Conti for helpful discussions. We are grateful to Laura Vettori, Isabella Pesce, Michael Pancher and the professionals at our institutional facilities for technical help. CIBIO Core Facilities are supported by the European Regional Development Fund (ERDF) 2014–2020. We thank FACS core facility at Aarhus University for technical assistance related to human muscle FACS. Work in the authors’ laboratories is supported by AFM/Téléthon - France (grant No. 23758 to S.B.), Telethon and Provincia autonoma di Trento - Italy (grant No. TCP13007 to S.B.), POR FESR Regione Lombardia (FORCE-4-CURE, grant No. 2526393 to Y.T.), Ministero della Salute (grant No. RF-2016-02362263 to Y.T.), GFB-ONLUS (grant No. PR-0394 to Y.T.), Italian Ministry of University and Research (grant No. 201742SBXA to PB), Telethon Foundation (grant No. GGP19229 to P.B. and GMR23T1221 to S.B.), Novo Nordic Foundation (grant No. NNF17OC0027242 to N.J. and J.F.). S.M. was supported by Marie-Curie MSCA Global Fellowship (grant No. 101149789).

## AUTHOR CONTRIBUTIONS

Conceptualization: P.B., Y.T., L.G., K.V., N.J, J.F. and S.B.; Methodology: E.K., M.L., S.M., M.V., L.G., P.B., Y.T., J.B.J., J.W. J.F. and S.B.; Validation: E.K., M.L. and S.B.; Formal Analysis: E.K., M.L., F.M., M.V., L.G., J.B.J., J.W. J.F. and S.B.; Investigation (all experiments): E.K. and M.L.; Investigation (Col6 experiments): M.L., S.D.S., and S.M.; Investigation (lineage experiments): E.K. and F.M.; Investigation (human cells isolation and characterization): M.L., M.B., J.B.J., J.W., N.J., J.F.; Investigation (bioinformatics analysis): M.V. and L.G.; Investigation (Single Cell RNAseq and CyTOF analysis): L.G.; Investigation (cellular and histological analysis): E.K., M.L., F.F., J.B.J., J.W., S.D.S., J.F. and S.B.; Resources: P.B, Y.T., C.V., N.J., J.F. and S.B.; Data Curation: M.L., E.K., J.B.J., J.W. J.F. and S.B.; Writing, Reviewing and Editing the manuscript: E.K., M.L., P.B., K.V., Y.T., L.G., J.F. and S.B.; Visualization: E.K., M.L., F.M., F.F. and S.B.; Supervision: J.F. and S.B.; Project Administration: E.K., M.L., N.J., J.F., Y.T. and S.B.; Funding Acquisition: Y.T., N.J., J.F. and S.B.

## RESOURCE AVAILABILITY

### Lead contact

Further information and requests for resources and reagents should be directed to and will be fulfilled by the lead contact, Stefano Biressi.

### Materials Availability

This study did not generate any unique reagents.

### Data and code availability

The data generated in this study are provided in figures and supplemental information. Row data are available upon request.

