## Supplemental Files for "CD90 identifies distinct fractions of muscle stem cells with different modalities of activation and quiescence maintenance"

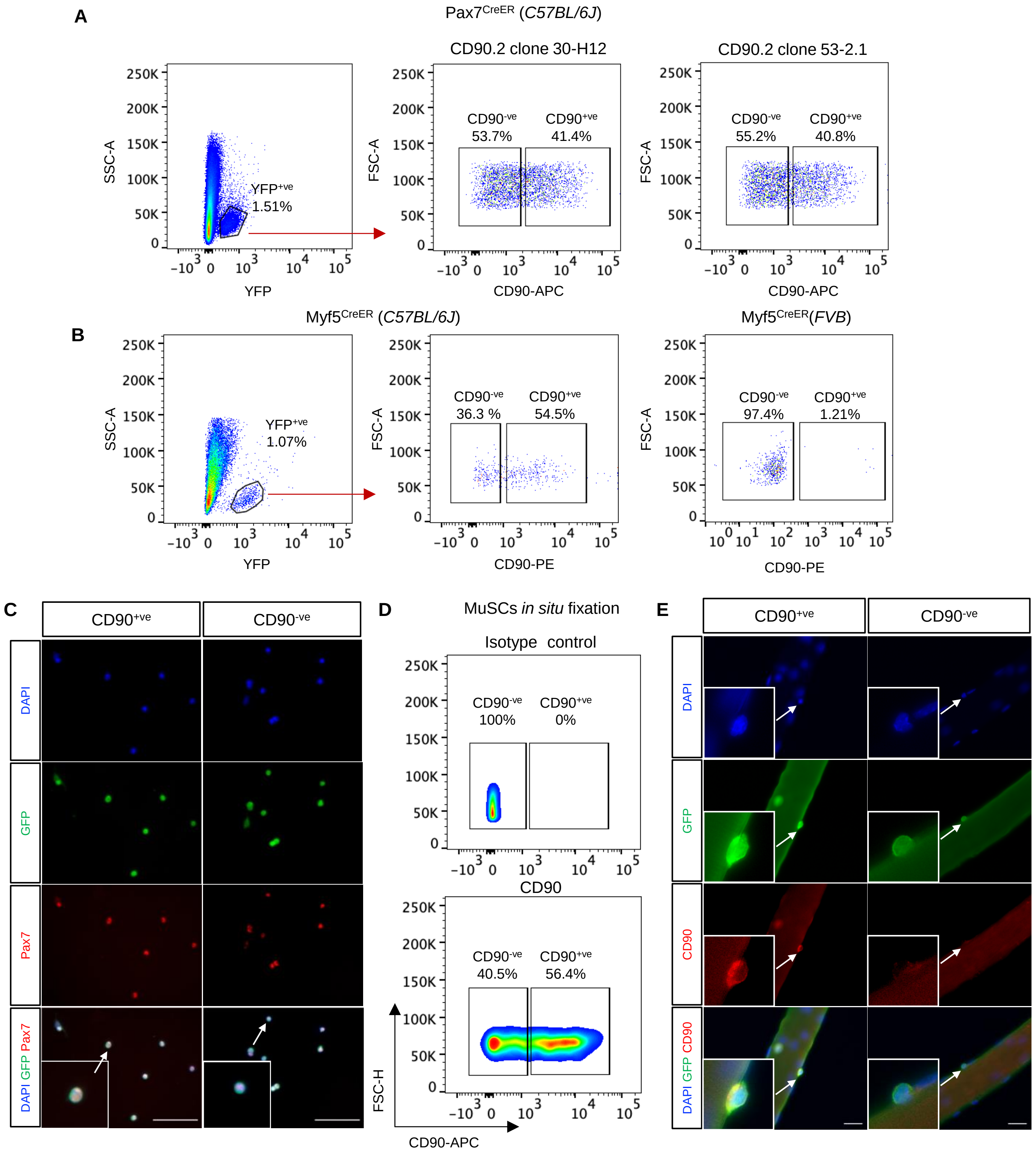


Figure S1 (related to Figure 1)

**Figure S1. Identification of two novel subpopulations of MuSCs**, **related to Fig.1.**

(**A**) FACS profile of MuSCs from uninjured hindlimb muscles of *Pax7*$^{CreERT/wt2}$*;R26R*$^{EYFP/wt}$ mice in the *C57BL/6J* background stained with two different antibodies against CD90.2 (clones 30-H12 or 53-2.1).

(**B**) FACS profile of MuSCs from uninjured hindlimb muscles of *Myf5*$^{CreER/wt}$*;R26R*$^{EYFP/wt}$ mice in the *C57BL/6J* (left) or *FVB* background (right) stained with an anti-CD90.2 antibody (clone 53-2.1).

(**C**) Immunofluorescence staining of Pax7 and YFP in CD90$^{+ve}$ and CD90$^{-ve}$ MuSCs isolated from uninjured hindlimb muscles of *Pax7*$^{CreERT2/wt}$*;R26R*$^{EYFP/wt}$ mice, and let adhere to the wells for 2 hours. DAPI was used to stain the nuclei. White arrows indicate magnifications of the CD90$^{+ve}$ or CD90$^{-ve}$ MuSCs (insets). Scale bar: 50 μm.

(**D**) FACS profile of *in situ* fixed YFP$^{+ve}$ MuSCs from *Pax7*$^{CreERT2/wt}$*;R26R*$^{EYFP/wt}$ hindlimb muscles stained with an isotype control (upper panel) and anti-CD90 (lower panel) antibodies.

(**E**) Freshly-isolated *Pax7*$^{CreERT2/wt}$*;R26R*$^{EYFP/wt}$ myofibers from *soleus* muscle, showing CD90$^{+ve}$ and CD90$^{-ve}$ MuSCs. White arrows indicate magnifications of the CD90$^{+ve}$ or CD90$^{-ve}$ MuSCs (insets). Scale bar: 20 μm.

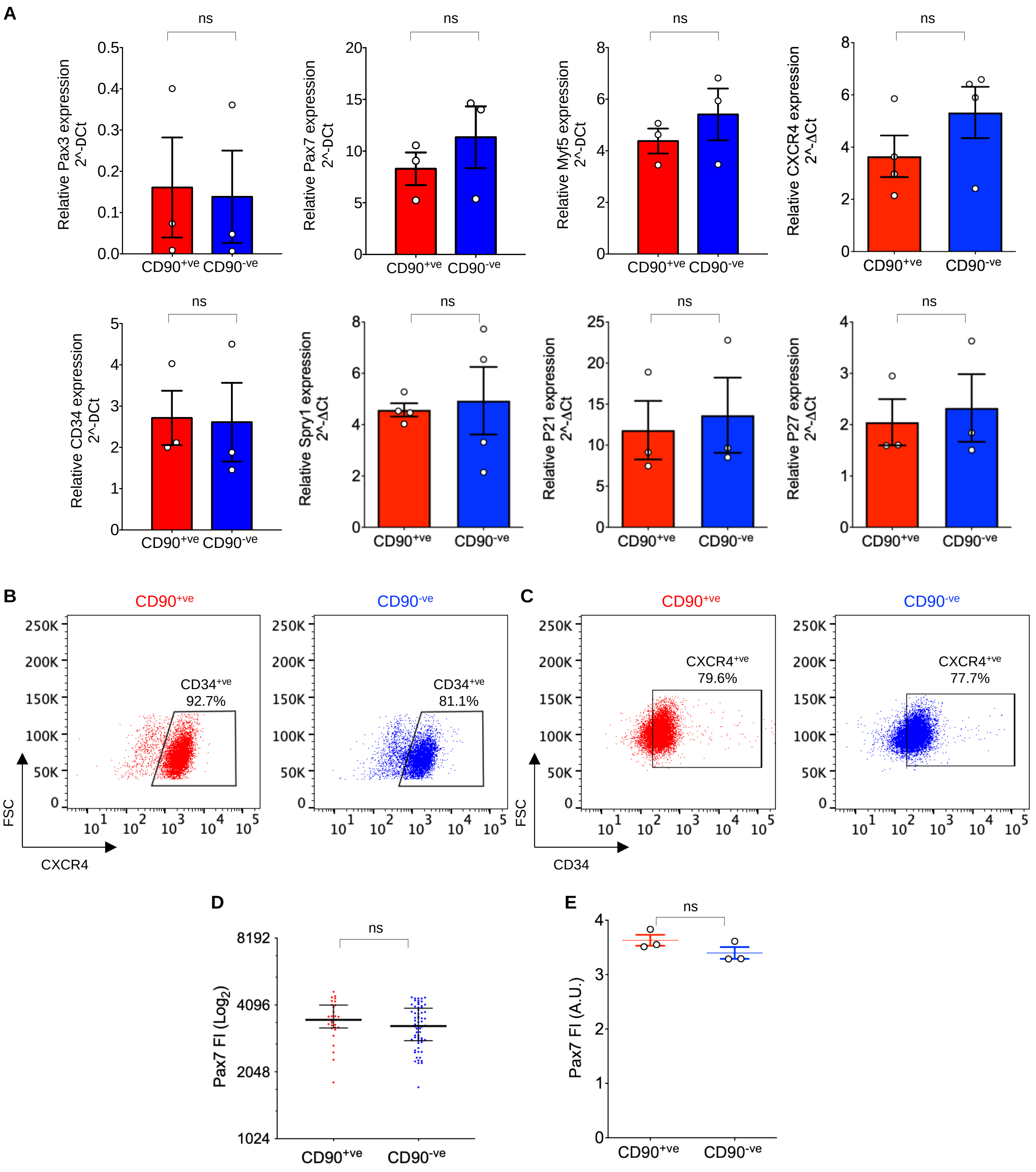


Figure S2 (related to Figure 1)

**Figure S2. Identification of two novel subpopulations of MuSCs, related to Fig.1.**

(**A**) RT-qPCR analysis of relative gene expression for known markers of MuSC heterogeneity in CD90$^{+ve}$ and CD90$^{-ve}$ subpopulations of MuSCs. Error bars represent mean ± SEM, n ≥ 3.

(**B-C**) FACS profile of CXCR4 (B) and CD34 (C) expression in CD90$^{+ve}$ or CD90$^{-ve}$ subpopulations of MuSCs from *Pax7$^{CreERT2/wt}$;R26R$^{EYFP/wt}$* (CXCR4) and wild-type (CD34) uninjured hindlimb muscles. Note that anti-CXCR4, anti-CD34 and anti-CD90 antibodies stain different fractions of cells.

(**D**) Example of single immunofluorescence evaluation of Pax7 fluorescence intensity (FI) in freshly isolated CD90$^{+ve}$ and CD90$^{-ve}$ MuSCs. Error bars represent the median with an interquartile range. A two-tailed Mann-Whitney test was performed.

(**E**) Pax7 fluorescence intensity (FI) in freshly isolated CD90$^{+ve}$ and CD90$^{-ve}$ MuSCs as quantified in (D). Error bars represent mean ± SEM, n = 3. A.U.: arbitrary units.

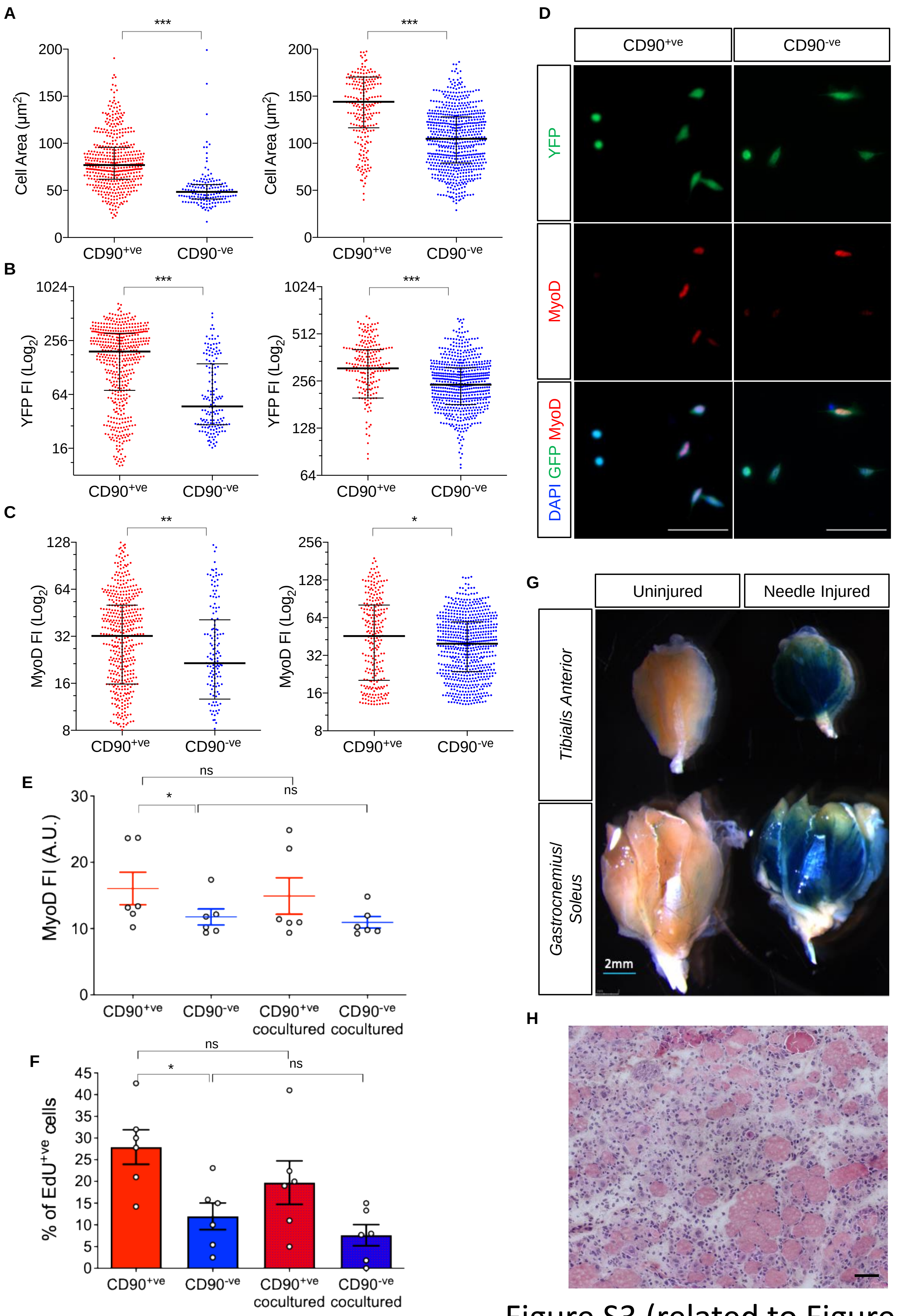


Figure S3 (related to Figure 2)

**Figure S3.** ***In vitro*** **and** ***in vivo*** **activation of MuSCs, related to Fig.2.**

(**A**-**C**) Additional measurements of cell area (A), YFP (*Rosa26* locus) (B), and MyoD fluorescence intensity (FI) (C), in FACS-isolated CD90$^{+ve}$ and CD90$^{-ve}$ MuSCs after 16 hours in culture (see Figure 2B-D). Error bars represent median with interquartile range, the number of cells is ≥ 150/replicate. Two-tailed Mann-Whitney test was performed.

(**D**) Representative immunofluorescence staining of MyoD and YFP in CD90$^{+ve}$ and CD90$^{-ve}$ MuSCs FACS-isolated from uninjured hindlimb muscles of $Pax7^{CreERT2/wt};R26R^{EYFP/wt}$ mice after 16 hours in culture. DAPI was used to stain the nuclei. Note the higher number of cells expressing MyoD above the background level in CD90$^{+ve}$ cultures compared to the CD90$^{-ve}$ counterpart. Scale bar: 20 μm.

(**E**-**F**) Quantification of MyoD fluorescence intensity (FI) (E) and percentage of EdU incorporating cells (F) in FACS-isolated CD90$^{+ve}$ and CD90$^{-ve}$ MuSCs maintained alone or co-cultured for 2.5 days *in vitro*. Note that the co-culture of CD90$^{+ve}$ and CD90$^{-ve}$ MuSCs does not affect the activation and proliferation of the single subpopulations. EdU was administered 2 hours before fixation. Error bars represent mean ± SEM, n=6. A.U.: arbitrary units.

(**G**) X-Gal staining of *tibialis anterior* (TA) and *gastrocnemius/soleus* muscles uninjured or needle-injured muscles from a $Pax7^{CreERT2/wt};R26R^{LacZ/wt}$ mouse. Regenerating fibers are stained in blue. Scale bar: 2 mm.

(**H**) Representative H&E staining of needle-injured *gastrocnemius* muscle sections at 4.5 days post-injury. Scale bar: 50 μm.

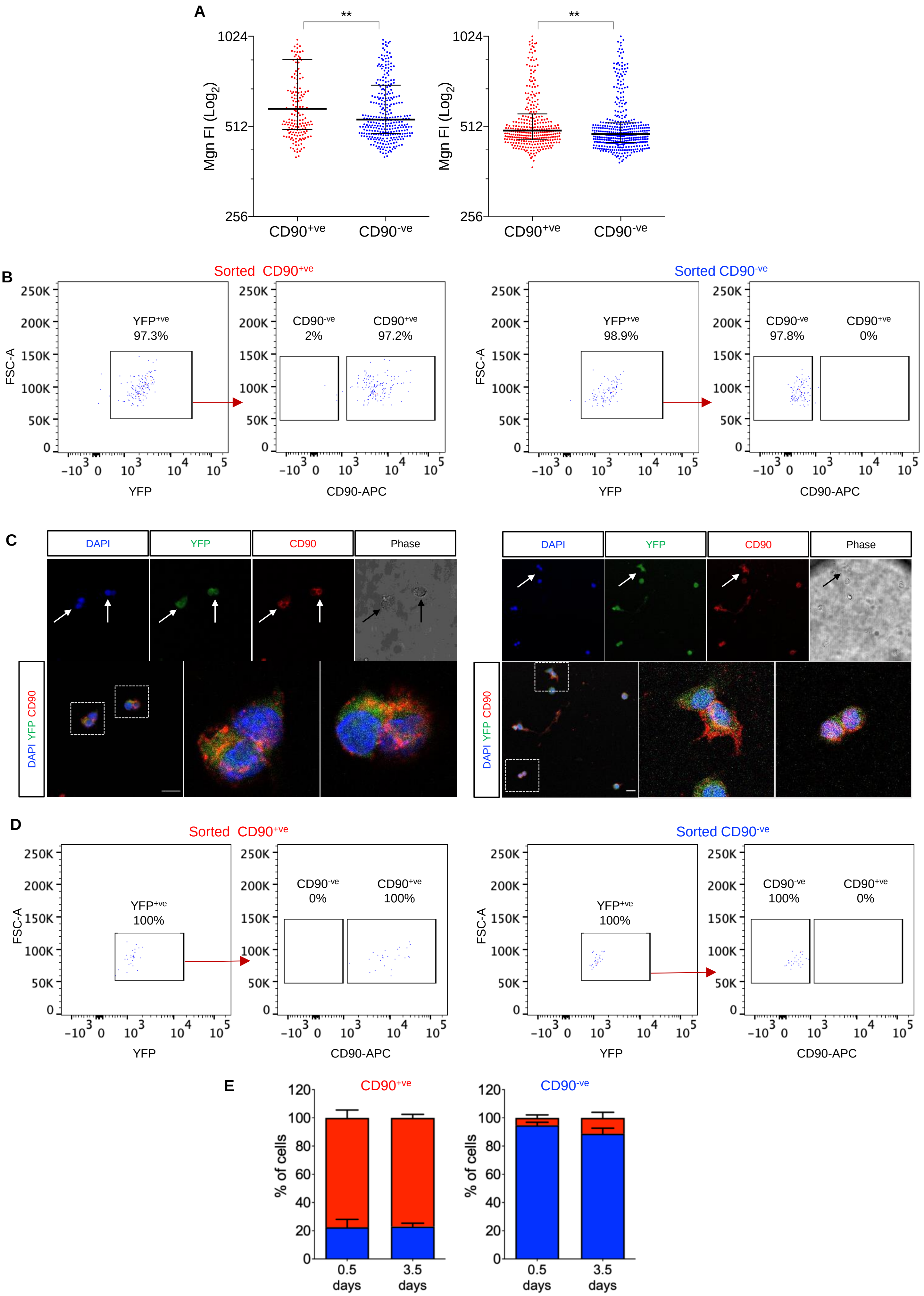


Figure S4 (related to Figure 3)

**Figure S4**. **Polarized expression of CD90, related to Fig. 3.**

(**A**) Additional measurements of Mgn fluorescence intensity (FI) of CD90$^{+ve}$ and CD90$^{-ve}$ MuSCs isolated 7 days post-injury (see Figure 3C). Error bars represent the median with an interquartile range; the number of cells is ≥ 180/replicate. A two-tailed Mann-Whitney test was performed.

**(B)** Representative FACS analysis of CD90$^{+ve}$ and CD90$^{-ve}$ MuSCs subpopulations sorted from uninjured tamoxifen-injected *Pax7*$^{CreERT2}$*;R26R*$^{EYFP}$ mice in experiments described in Figures 3D-F, and S4C. Note the purity of the sorted cells in terms of YFP expression (i.e., MuSCs identity) and CD90 positivity/negativity.

(**C**) Confocal images of cell doublets deriving from single CD90$^{+ve}$ MuSCs isolated from uninjured hindlimb muscles of *Pax7*$^{CreERT2/wt}$*;R26R*$^{EYFP/wt}$ mice. Arrows indicate newly formed YFP$^{+ve}$ MuSCs with polarized expression of CD90. Arrows indicate cells with asymmetric distribution of CD90. DAPI was used to stain the nuclei. Scale bar: 20 µm.

(**D**) Representative FACS analysis to assess purity of CD90$^{+ve}$ and CD90$^{-ve}$ MuSCs sorted subpopulations in experiments described in Figure S4E.

(**E**) Quantification of the percentage of CD90$^{+ve}$ and CD90$^{-ve}$ cells within the CD90$^{+ve}$ and CD90$^{-ve}$ populations sorted from uninjured tamoxifen-injected *Pax7*$^{CreERT2/wt}$*;R26R*$^{EYFP/wt}$ mice, plated in DMEM, 10% FBS, 2.5 ng/ml human FGF and analyzed at 0.5 and 3.5 days post-plating. Error bars represent mean ± SEM, n=3.

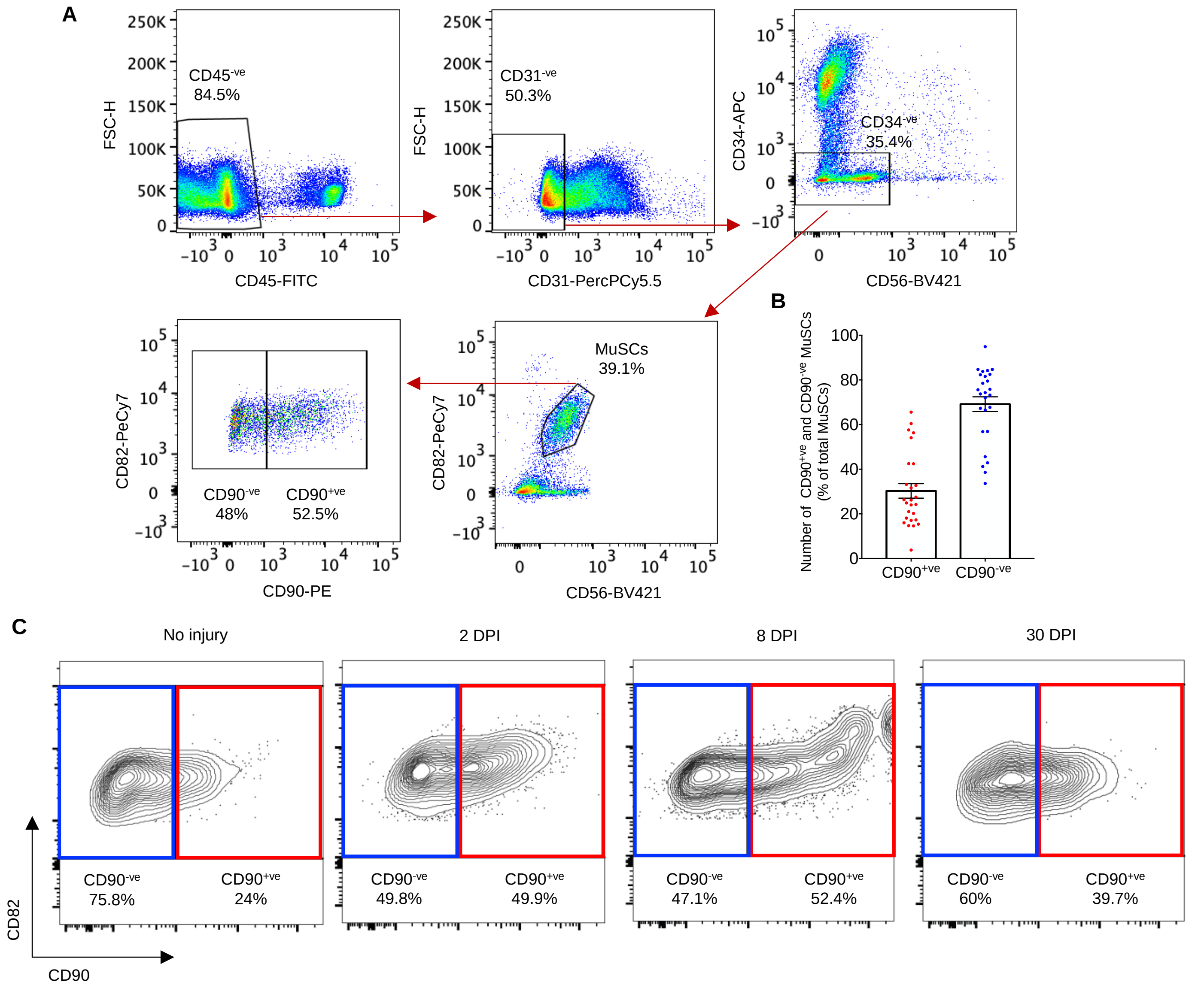


Figure S5 (related to Figure 4)

**Figure S5. CD90$^{+ve}$ and CD90$^{-ve}$ MuSCs diversification in human muscle, related to Fig. 4.**

(**A**) FACS profile of CD90 expression in MuSCs from human muscle negatively selected with CD45, CD31 and CD34 antibodies, and positively stained with CD82 and CD56 antibodies.

(**B**) Quantification of CD90$^{+ve}$ and CD90$^{-ve}$ MuSCs identified in human uninjured muscle as in (A). Error bars represent mean ± SEM, n=26.

(**C**) Example contour flow-plots displaying the distribution of CD90$^{+ve}$ and CD90$^{-ve}$ MuSCs in a 67-year old male before (-14) and 2, 8 and 30 days post injury (DPI). Biopsies were obtained from vastus lateralis muscle at all time-points. Note the pronounced expression of the activation determinant CD82 in CD90$^{+ve}$ MuSCs after injury.

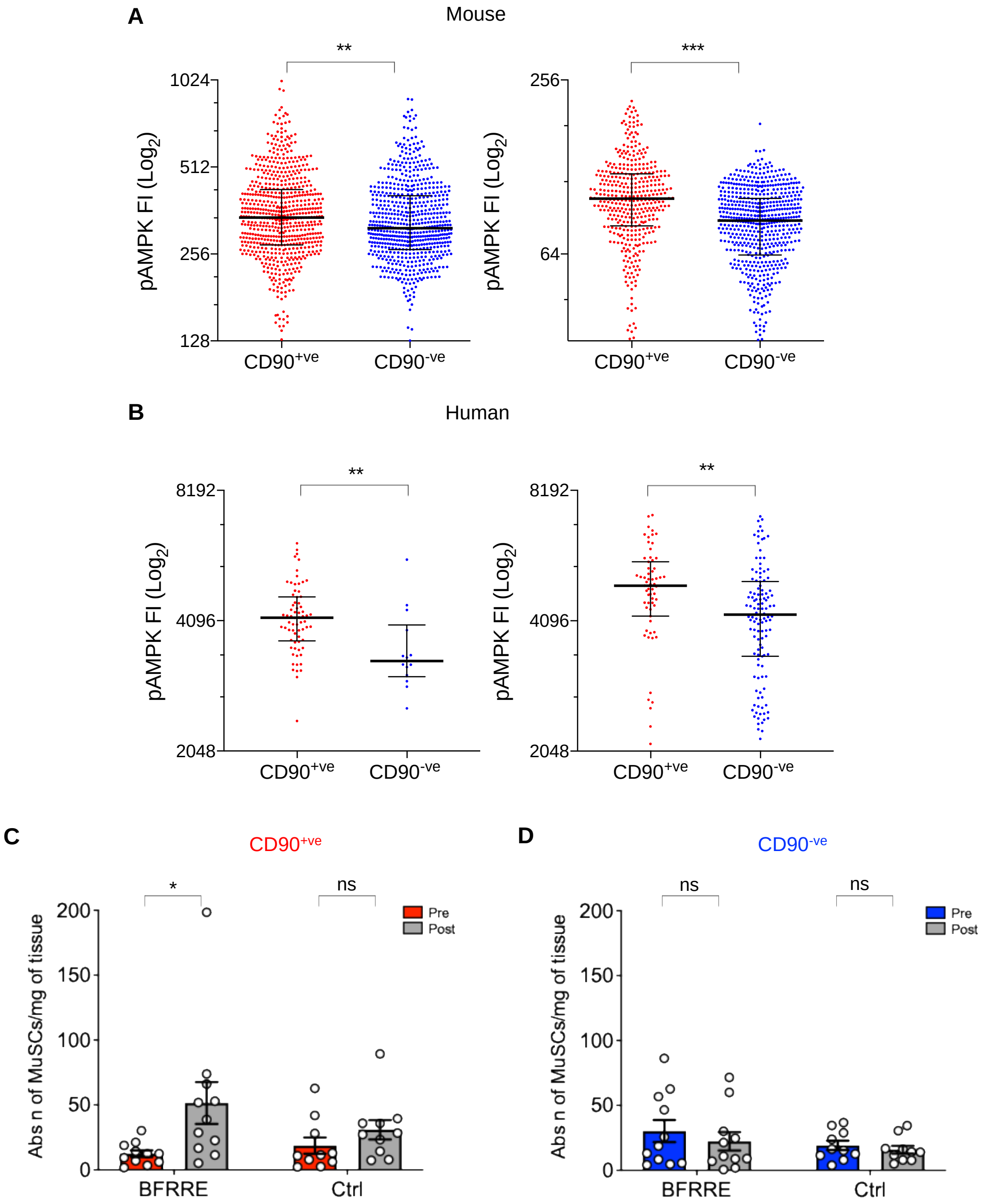


Figure S6 (related to Figure 5)

**Figure S6. CD90 is a determinant of functional heterogeneity, related to Fig. 5.**

(**A**, **B**) Additional replicates of $^{p}$AMPK fluorescence intensity (FI) analysis in mouse (A) and human (B) FACS-isolated CD90$^{+ve}$ and CD90$^{-ve}$ MuSCs (see Figure 5I and K). Error bars represent the median with an interquartile range. A two-tailed Mann-Whitney test was performed.

(**C-D**) Quantification of CD90$^{+ve}$ (C) and CD90$^{-ve}$ (D) MuSCs per mg of tissue in human muscles subjected to low-load blood flow-restricted resistance exercise (BFRRE) and their controls. Error bars represent mean ± SEM, n =11 (BFRRE) and 10 (Ctrl). A two-way Anova test was performed.

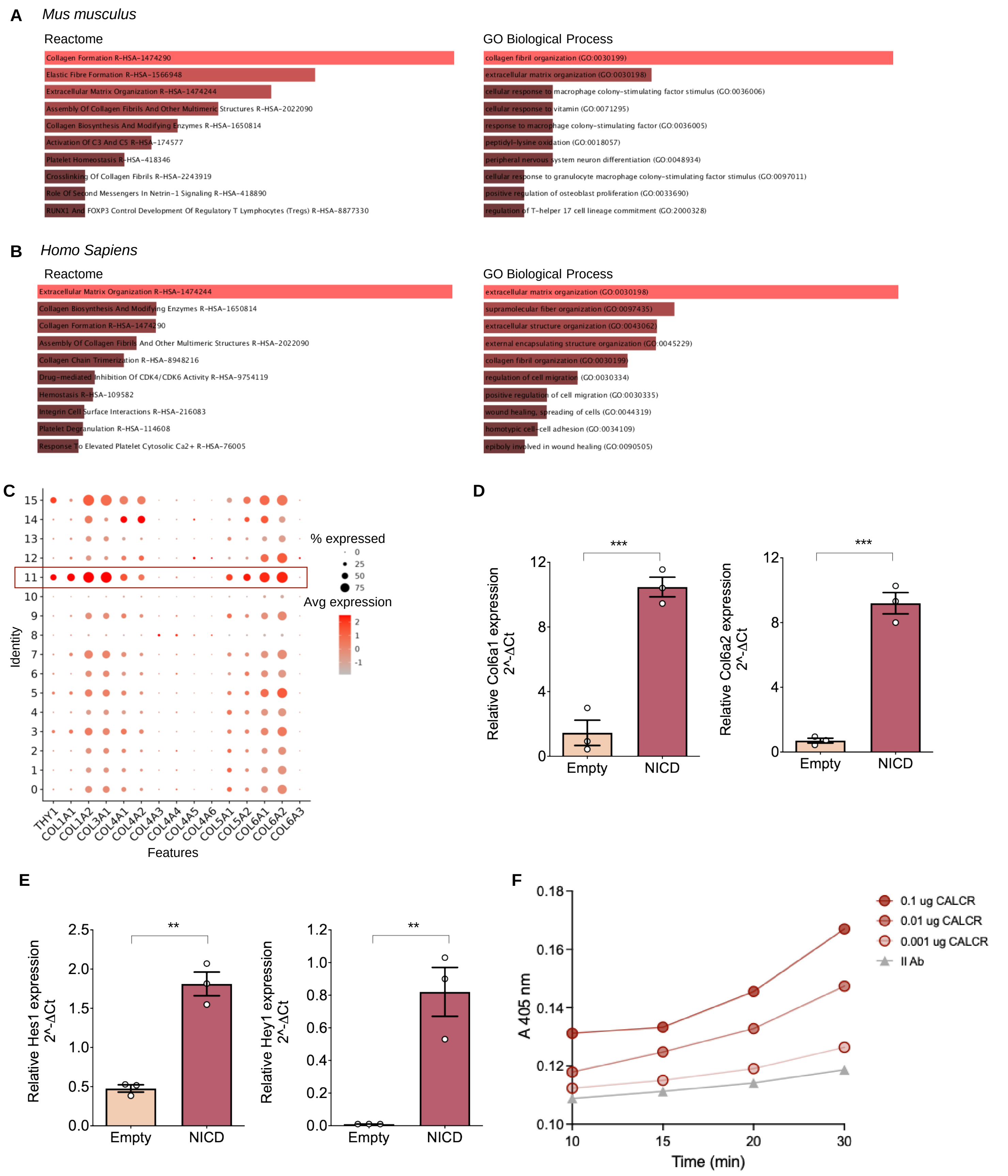


Figure S7 (related to Figure 6)

**Figure S7. CD90$^{+ve}$ MuSCs express ECM proteins, related to Fig. 6.**

(**A**, **B**) Reactome and Gene ontology (GO) analysis of differentially up-regulated genes in murine (A) and human (B) CD90-enriched MuSCs.

(**C**) Dot plot displaying the average expression and percentage of cells expressing selected ECM proteins in human CD90-enriched MuSCs (cluster 11).

(**D**) RT-qPCR analysis of *Col6a1* and *Col6a2* genes in C2C12 cells transfected with control (empty plasmid) or Notch intracellular domain (NICD)-expressing plasmids. Error bars represent mean ± SEM, n=3 biological replicates.

(**E**) RT-qPCR analysis of Notch target genes *Hes1* and *Hey1* genes in C2C12 cells transfected as in D. Error bars represent mean ± SEM, n=3 biological replicates.

(**F**) ELISA showing dose-response binding of CALCR to Col6 at specified time-points.

A

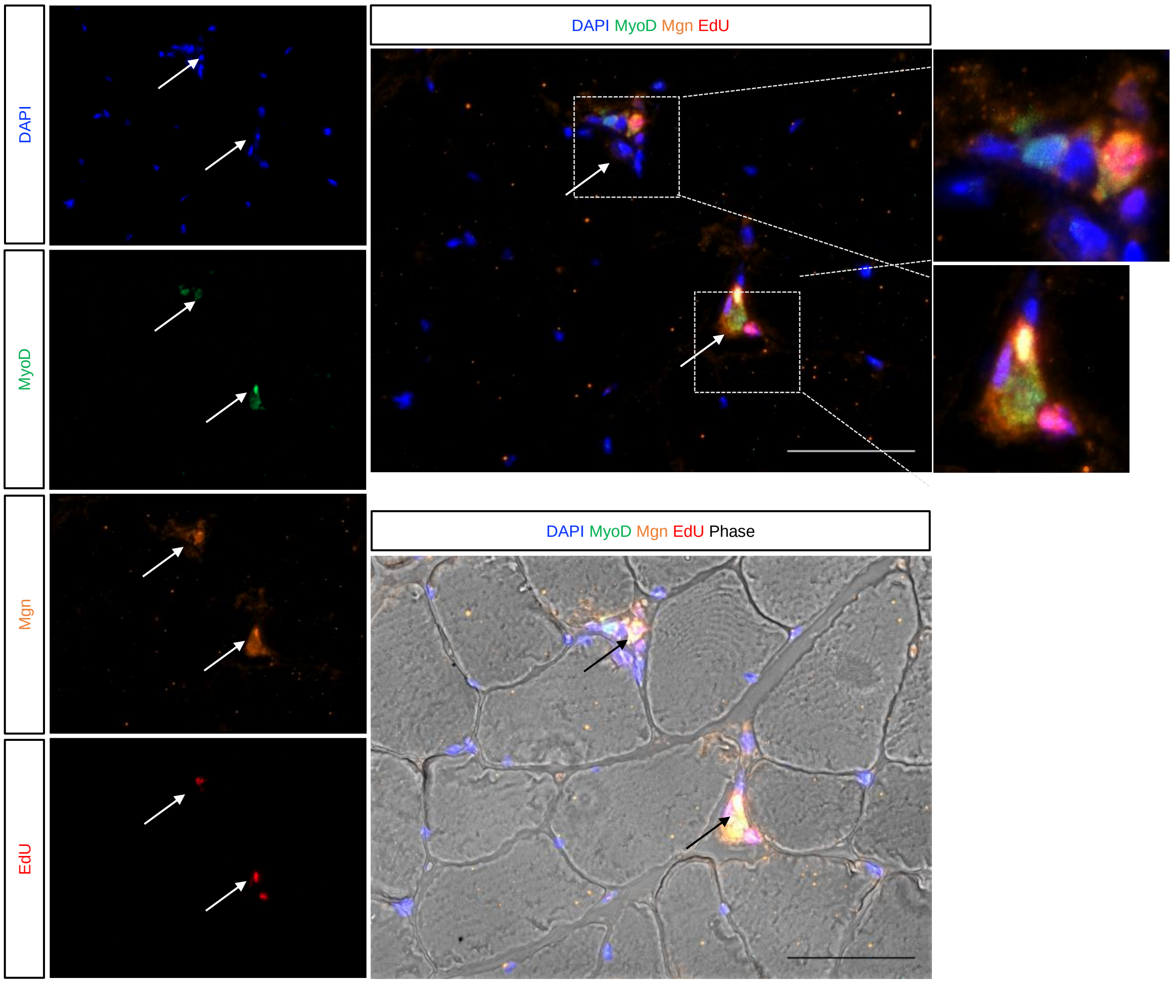


B

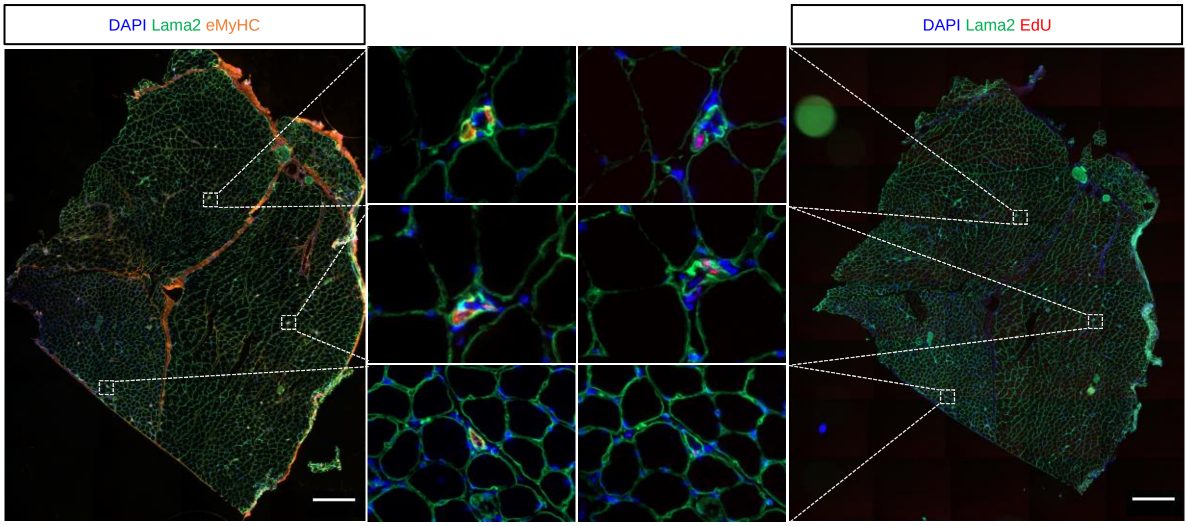


Figure S8 (related to Figure 6)

**Figure S8. MuSCs lineage progression in *Col6a1*$^{-/-}$ mice, related to Fig. 7.**

(**A**) *Gastrocnemius* muscle sections of uninjured *Col6a1*$^{-/-}$ mice injected with EdU for 5 consecutive days and stained with antibodies against MyoD and Mgn. Nuclei are visualized with DAPI. Arrows indicate Edu$^{+ve}$/MyoD$^{+ve}$/Mgn$^{+ve}$ cells resulting from MuSC activation and lineage progression. Scale bar: 50 μm.

(**B**) Serial *gastrocnemius* muscle sections of uninjured *Col6a1*$^{-/-}$ mice injected with EdU for 5 consecutive days and stained with Laminin α2 (Lama2) and embryonic Myosin (eMyHC) (left) or EdU (right). DAPI was used to stain the nuclei. Magnifications of the areas within dotted squares are shown in the center. Scale bar: 200 μm.

### SUPPLEMENTARY TABLES

#### Table S1. List of genes upregulated in human CD90-enriched MuSCs (cluster 11) with p-value <1.00E-53

| *Gene* | *Gene ID* | *Description* | *Adjusted p-value* |
|---|---|---|---|
| *COL3A1* | 1281 | Collagen type III alpha 1 chain | 0 |
| *COL1A2* | 1278 | Collagen type I alpha 2 chain | 0 |
| *COL1A1* | 1277 | Collagen type I alpha 1 chain | 1.10E-256 |
| *CAV1* | 857 | Caveolin 1 | 3.08E-248 |
| *GPC3* | 2719 | Glypican 3 | 9.91E-217 |
| *CD90/THY1* | 7070 | Thy-1 cell surface antigen | 6.77E-196 |
| *TWIST1* | 7291 | Twist family bHLH transcription factor 1 | 3.88E-182 |
| *LRRC17* | 10234 | Leucine rich repeat containing 17 | 2.53E-179 |
| *MDK* | 4192 | Midkine | 1.84E-172 |
| *COL6A2* | 1292 | Collagen type VI alpha 2 chain | 4.68E-165 |
| *CTHRC1* | 115908 | Collagen triple helix repeat containing 1 | 1.85E-158 |
| *MME* | 4311 | Membrane metalloendopeptidase | 4.18E-144 |
| *SPATS2L* | 26010 | Spermatogenesis associated serine rich 2 like | 1.28E-133 |
| *TNNT1* | 7138 | Troponin T1, slow skeletal type | 4.29E-132 |
| *COL6A1* | 1291 | Collagen type VI alpha 1 chain | 6.98E-131 |
| *CFH* | 3075 | Complement factor H | 2.40E-114 |
| *CD44* | 960 | CD44 molecule | 2.48E-101 |
| *NEAT1* | 283131 | Nuclear paraspeckle assembly transcript 1 | 4.39E-98 |
| *SESN3* | 143686 | Sestrin 3 | 2.70E-95 |
| *DKK3* | 27122 | Dickkopf WNT signaling pathway inhibitor 3 | 1.66E-92 |
| *CD82* | 3732 | CD82 molecule | 2.41E-92 |
| *PRSS23* | 11098 | Serine protease 23 | 6.09E-78 |
| *PCOLCE* | 5118 | Procollagen C-endopeptidase enhancer | 1.87E-77 |
| *RPL28* | 6158 | Ribosomal protein L28 | 4.15E-75 |
| *SPARC* | 6678 | Secreted protein acidic and cysteine rich | 5.06E-75 |
| *AEBP1* | 165 | AE binding protein 1 | 1.55E-71 |
| *MARCKS* | 4082 | Myristoylated alanine rich protein kinase C | 2.96E-71 |
| *RGS3* | 5998 | Regulator of G protein signaling 3 | 8.07E-71 |
| *TMSB10* | 9168 | Thymosin beta 10 | 1.27E-70 |
| *CCNI* | 10983 | Cyclin I | 6.41E-69 |
| *TGFB1* | 7040 | Transforming growth factor beta 1 | 7.25E-69 |
| *LMO7* | 4008 | LIM domain 7 | 8.72E-69 |
| *THBS1* | 7057 | Thrombospondin 1 | 1.32E-67 |
| *TP53I3* | 9540 | Tumor protein p53 inducible protein 3 | 4.09E-64 |
| *MYL6B* | 140465 | Myosin light chain 6B | 1.49E-63 |
| *PCDH7* | 5099 | Protocadherin 7 | 1.94E-63 |
| *SERPINF1* | 5176 | Serpin family F member 1 | 8.46E-63 |
| *CCND3* | 896 | Cyclin D3 | 5.53E-60 |
| *CHRNA1* | 1134 | Cholinergic receptor nicotinic alpha 1 subunit | 5.51E-56 |
| *COL5A2* | 1290 | Collagen type V alpha 2 chain | 1.06E-54 |
| *ANKRD28* | 23243 | Ankyrin repeat domain 28 | 9.08E-54 |

### Table S2. List of FACS antibodies

| *Antibody* | *Clone* | *Source* | *Identifier* | *Concentration* |
|---|---|---|---|---|
| Mouse anti-CD29-FITC | K20 | Beckman Coulter | Cat#IMO791U | 1:30 |
| Rat anti-CD31-APC/Fire | 390 | BioLegend | Cat#102434 | 1:100 |
| Rat anti-CD31-FITC | 390 | eBioscience | Cat#11-0311-85 | 1:250 |
| Mouse anti-CD31-PE | WM59 | BD Biosciences | Cat#555446 | 1:15 |
| REAffinity anti-CD31-PercPVio 700 | REA730 | Miltenyi | Cat#130-110-673 | 2 µl/sample |
| Mouse anti-CD34-APC | 581 | BD Biosciences | Cat#555824 | 20 µl/sample |
| Rat anti-CD34-FITC | RAM34 | eBioscience | Cat#BMS11-0341-85 | 1:100 |
| Mouse anti-CD34-PE | AC136 | Miltenyi | Cat#130-081-002 | 1:30 |
| Rat anti-CD45-APC/Fire | 30-F11 | BioLegend | Cat#103154 | 1:100 |
| Rat anti-CD45-FITC | 30-F11 | BioLegend | Cat#103108 | 1:250 |
| Mouse anti-CD45-FITC | 5B1 | Miltenyi | Cat#130-114-567 | 12 µl/sample |
| Mouse anti-CD45-PE | F10-89-4 | Southern Biotech | Cat#9625-09 | 1:300 |
| Mouse anti-CD56-BV786 | NCAM16.2 | BD Biosciences | Cat#564058 | 1:100 |
| Mouse anti-CD56-BV421 | NCAM16.2 | BD Biosciences | Cat#562752 | 1:100 |
| REAffinity anti-CD82-PE-Vio 700 | REA221 | Miltenyi | Cat#130-101-302 | 10 µl/sample |
| Rat anti-CD90.2-APC | 53-2.1 | BioLegend | Cat#140312 | 1:100 |
| Rat anti-CD90.2-APC | 30-H12 | BioLegend | Cat#105311 | 1:100 |
| Rat anti-CD90.2-BV421 | 53-2.1 | BioLegend | Cat#105341 | 1:100 |
| Mouse anti-CD90.2-PercP/Cy5.5 | 5E10 | BD Biosciences | Cat#561557 | 1:100 |
| Mouse anti-CD90-PE | 5E10 | Invitrogen | Cat#A15794 | 3.2 µl/sample |
| Rat anti-CD106 (VCAM)-Biotin | 429 | BioLegend | Cat#105704 | 1:100 |
| Mouse anti-CD184 (CXCR4)-APC | 12G5 | BD Biosciences | Cat#555976 | 1:15 |
| Rat anti-CD184 (CXCR4)-PercP/Cy5.5 | L276F12 | BioLegend | Cat#146509 | 1:100 |
| Rabbit anti-CALCR | - | BioRad | Cat#AHP3115 | 1:1000 |
| Rat anti-Ly6A/E (Sca-1)-BV421 | D7 | BioLegend | Cat#108127 | 1:100 |
| Rat anti-Ly6A/E (Sca-1)-PE | D7 | BioLegend | Cat#108126 | 1:100 |
| Streptavidin-APC | - | BioLegend | Cat#405207 | 1:100 |

### Table S3. List of human specimens used for flow cytometry analysis and cell sorting

| *Gender* | *Age* | *Experiment* |
|---|---|---|
| M | 69 | Fig. 4 A-B |
| M | 60 | Fig. 4B |
| F | 42 | Fig. 4B |
| F | 75 | Fig. 4F |
| M | 48 | Fig. 4F |
| F | 69 | Fig. 4F |
| M | 57 | Fig. 4F |
| M | 77 | Fig. 4F |
| F | 83 | Fig. 4F |
| F | 59 | Fig. S5B |
| F | 72 | Fig. 4H-J, 5A-B and S5B |
| F | 57 | Fig. S5B |
| M | 58 | Fig. S5B |
| M | 71 | Fig. 4H-J, 5A-B and S5B |

| | | |
|---|---|---|
| M | 56 | Fig. 4H-J, 5A-B and S5B |
| M | 70 | Fig. 4H-J, 5A-B and S5B |
| F | 67 | Fig. 4H-J, 5A-B and S5B |
| M | 67 | Fig. 4H-J, 5A-B and S5B and Fig. S5C |
| M | 62 | Fig. 4H-J, 5A-B and S5B |
| M | 61 | Fig. 4H-J, 5A-B and S5B |
| F | 63 | Fig. 4H-J, 5A-B and S5B |
| M | 70 | Fig. S5A |
| M | 75 | Fig. S5B |
| M | 75 | Fig. S5B |
| M | 69 | Fig. S5B |
| F | 63 | Fig. S5B |
| M | 61 | Fig. S5B |
| F | 60 | Fig. S5B |
| M | 70 | Fig. S5B |
| F | 63 | Fig. S5B |
| F | 78 | Fig. S5B |
| M | 75 | Fig. S5B |
| F | 71 | Fig. S5B |
| M | 79 | Fig. S5B |
| M | 66 | Fig. S5B |
| F | 68 | Fig. S5B |
| M | 63 | Fig. S6C-D |
| M | 66 | Fig. S6C-D |
| M | 56 | Fig. S6C-D |
| F | 57 | Fig. S6C-D |
| M | 72 | Fig. S6C-D |
| M | 66 | Fig. S6C-D |
| M | 65 | Fig. S6C-D |
| M | 72 | Fig. S6C-D |
| M | 75 | Fig. S6C-D |
| M | 67 | Fig. S6C-D |
| M | 67 | Fig. S6C-D |
| M | 72 | Fig. S6C-D |
| M | 59 | Fig. S6C-D |
| M | 62 | Fig. S6C-D |
| M | 60 | Fig. S6C-D |
| M | 68 | Fig. S6C-D |
| M | 63 | Fig. S6C-D |
| M | 73 | Fig. S6C-D |
| M | 60 | Fig. S6C-D |
| M | 64 | Fig. S6C-D |
| M | 68 | Fig. S6C-D |

### Table S4. Quantitative RT-PCR primers

| *Gene* | *Forward* | *Reverse* |
|---|---|---|
| CD34 | 5'-CTGACTTGAGAAAGCTGGGGA-3' | 5'-AAGATGGCCAGCAGAACTCC-3' |
| *Thy1* (CD90) | 5'-GAGTCCAGAATCCAAGTCGG-3' | 5'-CAAGACTGAGAGCAGGAGAGC-3' |
| *Calcr* | 5'-CGGCGGGATCCTATAAGTTG-3' | 5'-ATAGATCCCCTCGCAGAGCA-3' |
| *CD34* | 5'-CTGACTTGAGAAAGCTGGGGA--3' | 5'-AAGATGGCCAGCAGAACTCC-3' |
| *CDKN1A (P21)* | 5'-TGGAGTCAGGCGCAGATCCAC -3' | 5'- CGCCATGAGCGGCATCGCAATC-3' |
| *CDKN1B (P27)* | 5'-AGGCAAACTCTGAGGACCGGCA -3' | 5'- TGCTCCACAGTGCCAGCGTTC-3' |
| *Col6a1* | 5'-ACTGATGGGTGAGAGAGGTGA-3' | 5'-GCCTCTGTTGCCTGGATACC-3' |
| *Col6a2* | 5'-AAGACGTCCTTTGTCCAGACCC-3' | 5'-TGTGCACTGGGCCACATAGAG-3' |
| *Col6a3* | 5'-CCACTACTAAGCCCTTGGCGAG-3' | 5'-CTGGCACTGTTCTCCGTGAC-3' |
| *CXCR4* | 5'-CTTGACTGGCATAGTCGGCA-3' | 5'-TGTCCGTCATGCTCCTTAGC-3' |
| *Hes1* | 5'-GCCTCTGAGCACAGAAAGTCAT-3' | 5'-TCAGTGTTTTCAGTTGGCTTAGAC-3' |
| *Hey1* | 5'-AATGGAAACTTGAGTTCGGCG-3' | 5'-CTTCTCGATGATGCTCTCC-3' |
| *Hprt* | 5'-GGTTAAGCAGTACAGCCCCA-3' | 5'-CAAATCCAACAAAGTCTGGCCT-3' |
| *Myf5* | 5'-CACCTCCAACTGCTCTGACG-3' | 5'-AGCACATGCATTTGATACATCAGG-3' |
| *Myog* | 5'-TCCCAACCCAGGAGATCATTTG-3' | 5'-GTTGGGCATGGTTTCGTCTG-3' |
| *Pax3* | 5'-TTCCCGCTGGAAGTGTCC-3' | 5'-CAGAGGCCTGCCGTTGATAA-3' |
| *Pax7* | 5'-CTGGAAGTGTCCACCCCTCT-3' | 5'-TTGTGACGGATGTGGTTCGG-3' |
| *Sprouty 1 (Spry1)* | 5'-GAGGCCGAGGATTTCAGATGCA-3' | 5'-CTGAATCACCACTAGCGAAGTGT-3' |

### Table S5. List of primary antibodies used for immunofluorescence

| *Antibody* | *Clone* | *Source* | *Identifier* | *Concentration* |
|---|---|---|---|---|
| Chicken anti-GFP | - | Aves | Cat#1020 | 1:500 |
| Guinea pig anti-Col6 | - | - | Gara et al., 2011 | 1:300 |
| Mouse anti-MyoD1 | 5.8A | Dako | Cat#M351201-2 | 1:100 |
| Mouse anti-eMyHC | F1.652 | DSHB | Cat#F1.652 | 1:20 |
| Mouse anti-Mgn | 556358 | BD Biosciences | Cat#556358 | 1:100 |
| Mouse anti-Pax7 | - | DSHB | Cat#AB_528428 | 1:100 |
| Rabbit anti-CALCR | - | Biorad | Cat#AHP3115 | 1:100 |
| Rabbit anti-GFP | - | Invitrogen | Cat#A-11122 | 1:500 |
| Rabbit anti-pAMPK | - | Invitrogen | Cat#44-1150G | 1:250 |
| Rat anti-CD90.2-APC | 30-H10 | BioLegend | Cat#105311 | 1:100 |
| Rat anti-Laminin alpha 2 | 4H8-2 | Abcam | Cat#AB11576 | 1:1000 |

### Table S6. List of secondary antibodies used for immunofluorescence

| *Antibody* | *Source* | *Identifier* | *Concentration* |
|---|---|---|---|
| Chicken anti-Rat IgG (H+L), Alexa Fluor 647 | Invitrogen | Cat#A-21472 | 1:1000 |
| Donkey anti-Chicken IgG (H+L), Alexa Fluor 488 | EuroClone | Cat#J I703545155 | 1:1000 |
| Donkey anti-Mouse IgG (H+L), Alexa Fluor 594 | Invitrogen | Cat# A-21203 | 1:1000 |
| Donkey anti-Rabbit IgG (H+L), Alexa Fluor 488 | Invitrogen | Cat# A-21206 | 1:1000 |
| Donkey anti-Rabbit IgG (H+L), Alexa Fluor 594 | Invitrogen | Cat# A-21207 | 1:1000 |
| Donkey anti-Rat IgG (H+L), Alexa Fluor 488 | Invitrogen | Cat# A-21208 | 1:1000 |
| Donkey anti-Rat IgG (H+L), Alexa Fluor 594 | Invitrogen | Cat# A-21209 | 1:1000 |

### Table S7. List of siRNA targeting CD90

| *Name* | *Target sequence* |
|---|---|
| siRNA J-041986-09 | 5'-GUUAGAACAUAAGGGCGUA-3' |
| siRNA J-041986-19 | 5'-GGUCAAGUGUGGCGGCAUA-3' |
| siRNA J-041986-11 | 5'-GAGAGAAGAGGAAGCACGU-3' |
| siRNA J-041986-11 | 5'-GUAUCAGUGUGUAUAGAGA-3' |
